# Electroacupuncture at Governor Vessel improves neurobehavioral function via reduction of complexin1

**DOI:** 10.1101/787838

**Authors:** Yang Xu, Jia Liu, Xiaoming Zhao, Lei Zhou, Liuling Xiong, Cuiyun Li, Ya Jiang, Yangyang Wang, Tinghua Wang

## Abstract

Electroacupuncture at Governor Vessel (GV), as a traditional chinese medicine, has been proved that it can reduce scar and promote axon regeneration. However, the underlying mechanism remains unclear. Herein, complexin1 (CPLX1), as a candidate protein, was found using protein chip. Therefore, using a CRISPR/Cas9 knockout approach, we deleted CPLX1 specifically in the SD rats to assess the role of CPLX1 in GV treatment. Additionally, eIF5A1 stimulate the translation of CPLX1 with PPG sequence, we attempt to uncover whether eIF5A1 play a role in the GV treatment. In fact, GV can reduce scar and promote axon regeneration after SCC. CPLX1^−/+^ SCC rats demonstrated that decreased CPLX1 improved the microenvironment of injured area via reducing the components of fibrotic scar and further enhanced the synaptic plasticity, which benefit the regeneration of axons. And eIF5A1 could regulate the expression of CPLX1 in the process of GV treatment. Therefore, GV contributes to axon regeneration and synapse plasticity via eIF5A1 regulating CPLX1 following SCC, providing a convincible mechanism for improving the therapeutic efficacy of GV for SCC.

## Introduction

Spinal cord contusion (SCC) is a traumatic spinal cord injury (SCI) that can disrupt the communication between supraspinal centers and spinal circuits producing clinically irreversible disability and result in much comorbidity (Moraud *et al.*, 2016), which is characterized by neurological deficits and motor dysfunction with the subsequent tissue damage due to demyelination of axons and cellular death (Wang *et al.*, 2011). Terribly, it is estimated that only half of affected patients regain supraspinal control of movements below the level of the lesion after SCI (Asboth *et al.*, 2018). SCI causes profound and persistent neurological deficits, many of which result from the interruption of axonal connectivity (Asboth *et al.*, 2018). On the other hand, scar formation, a long-lasting inflammatory response and myelin debris prevent the axon regeneration after SCI (Chen *et al.*, 2018; Ruschel&Bradke, 2018). Thus, the prevailing model posits the benefit of promoting axonal extension by blocking extrinsic inhibitors or enhancing intrinsic activators when synapse exhibits the highest potential for plasticity (Wang *et al.*, 2011). Nowadays, numerous therapeutic strategies attempt to find the effective therapy eliminate these negative factors or promote axon regeneration, but there still is a lack of effective treatments for the SCC.

Governor Vessel electro-acupuncture(GV), as a therapeutic technique used in traditional Chinese medicine, is a type of therapy in which a needle inserted into an acupoint is attached to a trace pulse current with the purpose of producing synthetic electric and needling stimulation (Ding *et al.*, 2009). At present, GV has been applied profoundly in multiple fields. It has been reported that GV not only can induce the death of neural cells (Li *et al.*, 2012), but also improve median nerve function by somatotopically distinct neuroplasticity (Maeda *et al.*, 2017). In addition, many studies have proved that GV provides an effective strategy to alleviate motor dysfunction after SCI and this has resulted in promising functional recovery (Liu *et al.*, 2011; Li *et al.*, 2012; Zhang *et al.*, 2014; Liu&Wu, 2017). However, the mechanisms through which GV enables the production of motor patterns remain unclear, even though this understanding is pivotal in treating SCC. Given the satisfactory therapeutic effects, it was urgent to find the mechanism of GV on SCC.

CPLX1 is a presynaptic small molecule protein consisting of 134 amino acids, forming a SNARE complex with synaptobrevin, syntaxin, and snap 25 in the central nervous system, involving in anchoring, pre-excitation, and fusion of axonal end vesicles (McMahon *et al.*, 1995). Moreover, CPLX1 is highly homologous hydrophilic proteins that are tightly conserved, with 100% identity among mouse, rat, and human CPLX1. Nowadays, CPLX1 expressing has been widely studied in the disease related with neuroscience. Several lines of studies have linked CPLX1 to AD, demonstrating that CPLX1 was significantly down-regulated and the decline is positively correlated with the severity of AD (Nie *et al.*, 2017). Additionally, it was found that CPlX1 levels in the medial thalamus were decreased in Wernicke’s encephalopathy, suggesting that it may contribute to the pathophysiology of thalamic damage and offer a potential basis for the well-known differences in pathology between this structure and the inferior colliculus in this disorder (Hazell&Wang, 2005). Moreover, some researchers found that behavioural deficits in reversal learning were appeared in CPLX1 knockout mice as well Huntingdun mice, both showing abnormalities in long-term potentiation, which indicated that loss of CPLX1 is part of the underlying mechanism of the HD (Freeman&Morton, 2004; Glynn *et al.*, 2007). Therefore, these studies revealed that CPLX1 can play a role in the neuroscience. However, little effort has been focused on the motor function and the mechanism of CPLX1 remains unclear. Herein, we have addressed these important unresolved questions. We modeled spinal cord injuries in CPLX1 knock-out rats and analyzed the underlying mechanism of CPLX1 in SCC.

In this study, we reasoned that because the interruption of descending pathways abolishes the sources of modulation and excitation that are essential to enable functional states of spinal circuits, GV might stimulate axonal growth and recovery and enable individuals with motor paralysis to produce isolated leg movements after spinal injury. In order to test this hypothesis, we modeled these injuries in CPLX1 knock-out rats and utilized a GV treatment. Here, we provided evidence that GV promote the recovery of motor function via regulating CPLX1 using WB, EMG, MEP, DTI, Field Potential and Golgi staining. In addition, we attempted to prove that GV exert its role through eIF5A1 regulating CPLX1 using FM1-43 dyes *in vivo* and *in vitro*. Taken together, these data support a role for GV as a treatment for SCC and demonstrate the mechanism of GV treating SCC via eIF5A1 regulating CPLX1, establishing a reliable molecular theoretical basis for clinical medical workers in the treatment of spinal cord injury patients with GV.

## Results

### GV effectively improved the hindlimb motor dysfunction in SCC rats via increases synaptic plasticity through changing the axon components and transporter activity

As GV is a clinically feasible way to improve the motor dysfunction effectively, we aimed to decipher its distinct cellular actions. First, BBB score was performed to evaluate the effectiveness of GV on the motor function of SCC rats. Results indicated that the scores after GV increased obviously at 4 weeks post injury (wpi), suggesting that GV effectively improves hindlimbs motor dysfunction in SCC rats (Fig.1A, P<0.001). Moreover, compared to the SCC group, the GV-SCC groups showed significantly improvement in the MEP amplitude (Fig.2E-F, P<0.05), but, no significant difference existed in the latency (Fig.2E, G). Next, at gross anatomical levels, it was observed that GV treatment reduce the macroscopic scar area even SCC rats also display that the hematoma was disappearing gradually and scar was formed (Fig.1B, Fig.S1A). Interestingly, the images elicited by HE staining demonstrated the less formation of cavity and inflammatory cell infiltration after GV (Fig.1B). We next examined whether GV treatment might alleviate the collapse of micro-structures. Data of electron microscopy (EM) in Figure 1C showed significant protection of myelinated axons within the injured spinal cord of GV treated rats at 4 wpi (Fig.1C). Additionally, GV treatment significantly increased the percentage of presynaptic coverage of motor neurons after spinal cord injury (Fig.1C-D, P<0.05). These data suggested that GV treatment increase the axonal plasticity which was also confirmed via the synapsin I immunohistochemistry. Percentage area of synapsin-positive puncta caudal to the injury epicenter was used to quantify changes in synaptogenesis on ventral horn. There was a rising tendency of synaptic density with no statistical differences in the GV-SCC group when comparing to SCC group (Fig.4H-I, P>0.05). To acquire mechanistic insight, we performed protein mass spectrometry analysis of spinal cord segments including the injured site and regions immediately rostral and caudal to the epicenter over a period of 28 days after GV treatment. We identified 379 proteins that were differentially expressed including 226 up-regulated and 153 down-regulated proteins (false discovery rate [FDR]-adjusted p<0.05, fold change >2 or <0.5) (Fig.1E). Gene Ontology (GO) enrichment analysis of differentially expressed genes (DEGs) were conducted in three categories including biological process (BP), cellular component (CC) and molecular function (MF). Results demonstrated that almost DEGs locate in axon and synapse except extracellular space (Fig.1F). Transporter activity was the top enriched term in MF. While for BP, neurogenesis and neurotransmitter transport belonging to the top ten enriched term were focused (Fig.1F). Moreover, DEGs were selected again according the category of fold change >5 or <0.2 to further analysis these differentially expressed proteins. 23 up-regulated and 16 down-regulated proteins were picked (Fig.1G). Then, Venny 2.1.0 software was used to uncover the overlapping proteins which have striking differences and participate in focused MF and BP. A total of 5 up-regulated proteins (Slc12a5, Slc6a5, Slc6a11, Slc1a2 and Cplx1) were selected (Fig.1H).

**Figure 1.**
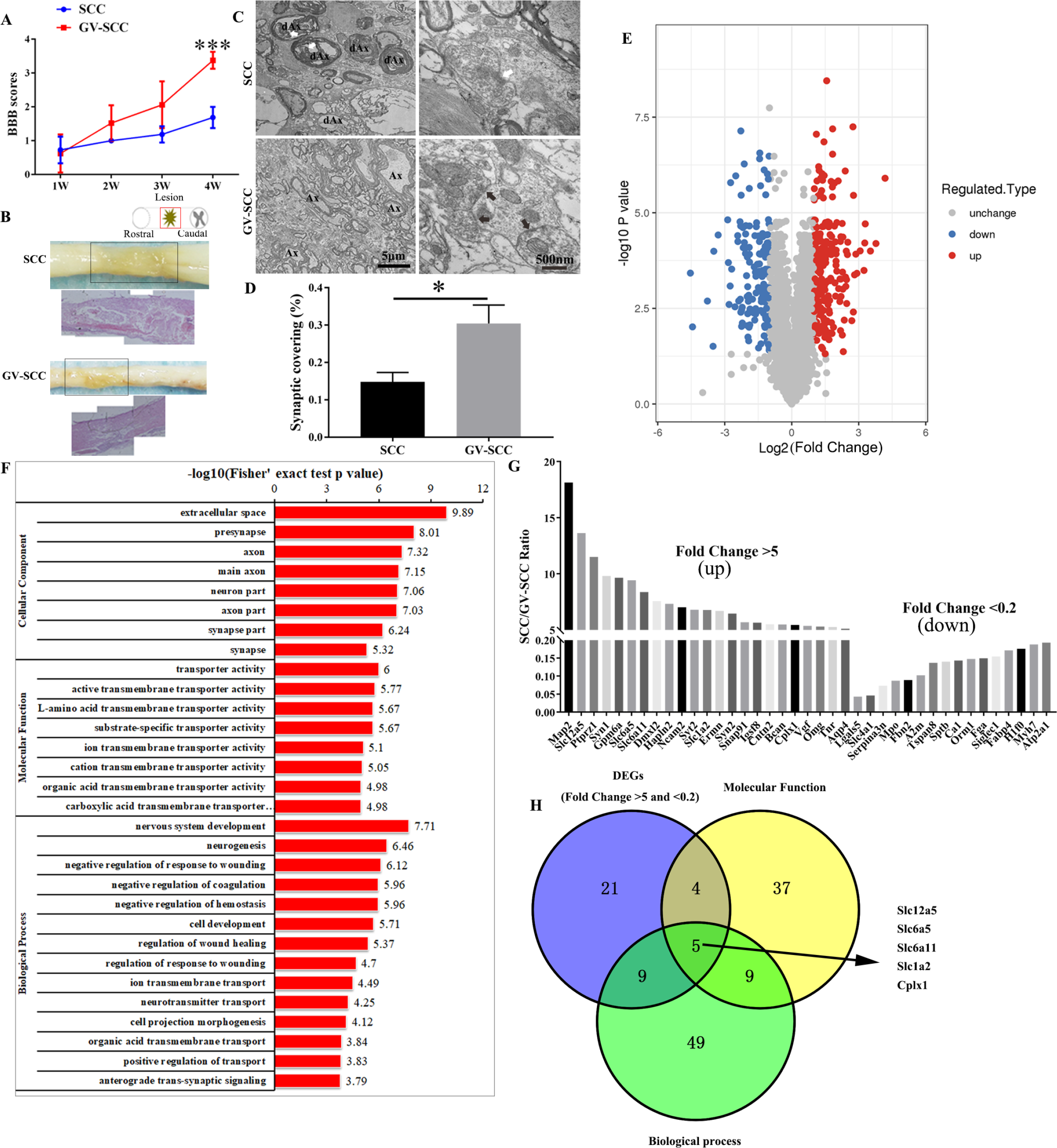
GV treatment accelerate the recovery of motor ability and increases synaptic plasticity after SCC which were related to down-regulating the expression of five candidates. A BBB scores of SCC and GV-SCC groups. B Low power photographs and sagittal view of the spinal cord of SCC and GV-SCC rats. Solid rectangles mark the injured area and the lower panels are the HE stain of lesion area. C Electron microscopic images of spinal cord sections rostral to the epicenter of the lesion in SCC rats and GV-treated rats n = 5 animals per group. D Percentage of motor neuron circumference covered by synapses within ventral horn. E Volcano plot of all differentially expressed genes (DEGs) between lesion sites of SCC and GV-SCC animals 4 wpi (weeks post injury). The data for all genes are plotted as log2 fold change of the adjusted p-value. F GO functional enrichment analysis of DEGs. G Bar graph showing the DEGs sorted according to the fold change >5 or <0.2. H Three-set Venn diagram represent the selected DEGs according to the indicated strategy. Scheme in (B) indicates lesion and displayed region (red box). wpi, weeks post injury. Data information: Data were analyzed using Student’s t-test. Values are plotted as means ± SEM. *P < 0.05, **P < 0.01, ***P < 0.001.

### Moderate reduction of CPLX1 could significantly improve the motor function for a relatively stable time after SCC even better than GV-treatment

It is well known that, excepting the scarring, axon growth inhibitory factors were the other key impediment to regenerating axons, in which chondroitin sulfate proteoglycans (CSPGs) is one of the major component (Ruschel *et al.*, 2013). Here, the effective function of five candidate proteins to promote the axon elongation were uncovered under an inhibitory environment in which CSPGs was added. Groups of slc6a11-siRNA and CPLX1-siRNA significantly restored axon growth when neurons growth in an inhibitory environment (Fig.2A-B, P<0.001 for both groups). And, down-regulation of CPLX1 showed even more breathtaking growth. Therefore, we next validate the role of CPLX1 in the GV treating SCC rats. As the axon trajectory of interneuron populations can vary and thus play different roles in different sites near a SCC, so we first determined whether neurons responded differently according to their location relative to the lesion site, and tissues in rostral, lesion and caudal sites were separated to do further analyze (Fig.1B). First, CPLX1 significantly increased in the rostral and caudal sites comparing with epicenters at different time points (Fig.S1C-D). The comparison assessed by WB in the same sites was made between the GV-SCC and SCC group demonstrating that the CPLX1 expression of rostral and caudal in the GV-SCC group decreased after GV comparing to SCC group (Fig.1J, P<0.05). Thus, these implied that silencing the expression of CPLX1 may be the effective mechanism of GV treatment.

**Figure 2.**
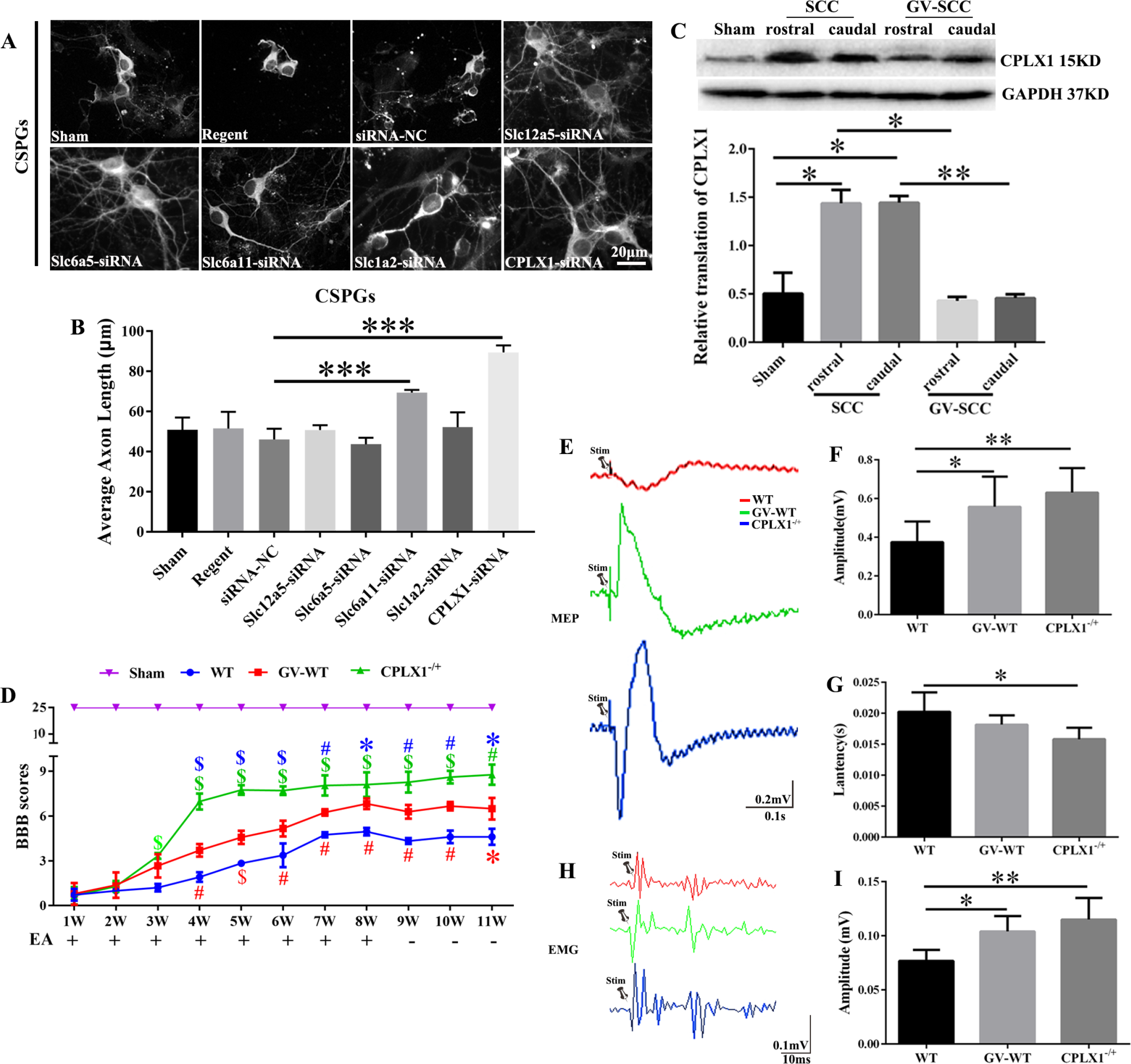
Moderate reduction of CPLX1 improve the MEP and EMG and enhances motor function of hind limbs simultaneously after SCC even more than GV application. A Beta-3 tubulin (Tuj-1) immunolabeling of neurons growing on inhibitory substrates (CSPGs, chondroitin sulfate proteoglycans) among different groups. B Quantification of neurite length of cortical neurons under different conditions, n=4 experiments. Values are plotted as means ± SEM. ***P < 0.001. C Western blots (WB) of CPLX1 in regions of rostral and caudal to the epicenter, n=3 animals per group. D BBB scores of the aforementioned experiments are shown, n=8 animals per group. Red symbols represent GV-WT group vs. WT group at corresponding time point; Green symbols represent CPLX1^−/+^ group vs. WT group at corresponding time point; Blue symbols represent CPLX1^−/+^ group vs. GV-WT group at corresponding time point; *P<0.05, ^#^P<0.01, ^$^P<0.001. E The figure shows representative MEP signal in different group, n=4 animals per group. F Quantification and statistical analyses of amplitude for the aforementioned experiments are shown. G Quantification of latency period for these groups. H Representative EMG signal in different group, n=4 animals per group. I Quantification of amplitude for these groups. Data information: Data were analyzed using one-way ANOVA. *P < 0.05, **P < 0.01, ***P < 0.001 in figure B, C, F, G and I. Values are plotted as means ± SEM.

For further researching, we used a transgenic rat line with homozygous knockout of CPLX1 generated by CRISPR/Cas9 gene editing system. Duo to CPLX1^−/−^ rats had a profound ataxia and short lifespan, CPLX1^−/+^ rats were used for the following experiment, which have been verified with significant levels of CPLX1 gene silencing (Fig.S1B, P<0.001 for both comparisons). Then, CPLX1^−/+^ and GV-WT group of SCC models were conducted for further investigating the role of CPLX1. Results assessed by BBB score suggested that CPLX1^−/+^ SCC rats got a better recovery than that of the WT SCC and GV-WT SCC group since the third week after injury, and this trend kept extending even at 9, 10, 11 week, on those period no GV carry out, indicating the functional sustainability in the GV treating and partial delection of CPLX1 in SCC rats with amazing outcome (Fig.2D). When the nervi ischiadicus was stimulated near the base of the tail in WT rats, little or no reflex was evoked in the segmental gastrocnemius muscle EMG recordings, regardless of the stimulation current intensity or rate of stimulation. However, the response was more obvious in the CPLX1^−/+^ and GV-WT group as shown in the results that the amplitudes of CPLX1^−/+^ group (P<0.01) and GV-WT (P<0.05) were larger than the WT group (Fig.2H-I). Besides, the MEP recordings of the WT rats remained stable and the evaluated measures showed steady and similar values for the duration of the testing period. The amplitude of CPLX1^−/+^ group (P<0.01) and GV-WT (P<0.05) was larger than the WT group. The latency in the CPLX1^−/+^ group was larger when compared to the WT group (P<0.05, Fig.2E-G).

### Moderate reduction of CPLX1 (CPLX1^−/+^) enhances neural tissue regeneration and reduces fibrotic scar tissue after SCC even more than GV application

As previously mentioned, reduction of CPLX1 using genetic method in the SCC models significantly ameliorated the motor dysfunction. Eleven weeks later, the content of CPLX1 at the rostral and caudal spinal cord extraction in the CPLX1^−/+^ SCC models was lower than that of WT SCC group, respectively (Fig.3A, P<0.05). Further, Diffusion tensor imaging (DTI) scans was used noninvasively to longitudinally track neural tissue regeneration progress, which was an ideal measurement for human SCI clinical studies. Results demonstrated that spinal cord neural regeneration was more obvious in CPLX1^−/+^ SCC rats. Different colors of tracking were used to mark the direction of fiber orientations. It was found that, in the WT rats, the ascending and descending blue fiber tracks were disconnected, leaving a gap on the center region of the spinal cord. While after GV, no gap existed in the spinal cord, and the blue fiber signals filled the whole spinal cord structure in the CPLX1^−/+^ rats. Quantitative analyses of percentage of rostral–caudal voxels and FA values indicated a higher value in CPLX1^−/+^ group compared with WT and GV-WT group, which demonstrated significant neural tissue regeneration in CPLX1^−/+^ rats and GV (Fig.3B).

**Figure 3.**
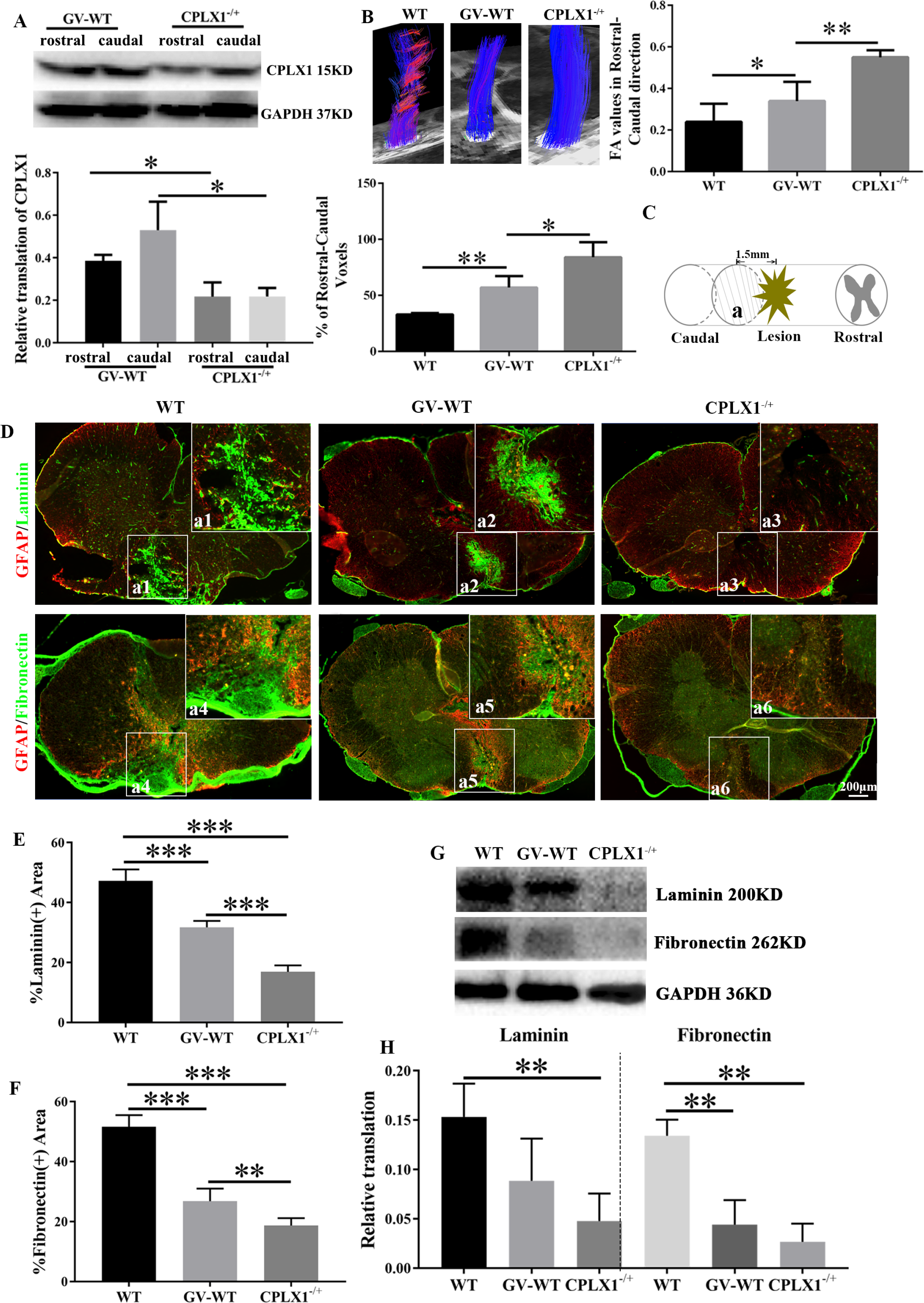
Moderate reduction of CPLX1 (CPLX1^−/+^) enhances neural tissue regeneration and reduces fibrotic scar tissue after SCC even more than GV application. A WB of CPLX1 in the candal and rostral spinal cord extracts among GV treatment group (GV-WT) and CPLX1^−/+^ group, n = 3 animals per group. B Typical fiber tract reconstruction for the wild-type SCC (WT), GV-WT and CPLX1^−/+^ groups is displayed. Graph of averaged FA values and percentages of rostral–caudal voxel numbers of the three groups in the area rostral and caudal 5mm to the lesion site. C Diagram illustrating the spinal cord contusion and displayed region (labeled as a) in the following figure 2D. D Immunolabeling of laminin, fibronectin and glial fibrillary acidic protein (GFAP) in transverse section in the indicated section marked in figure 2C (labeled as a) among these three groups. Scale bar, 20 μm. E, F GFAP- and Fibronectin-positive (+) area at the lesion site, n = 5 animals per group. G, H WB of laminin and fibronectin in caudal spinal cord extracts, n = 3 animals per group. Data information: Data were analyzed using one-way ANOVA. Values are plotted as means ± SEM. *P < 0.05, **P < 0.01, ***P < 0.001.

Function changes of caudal spinal cord to the lesion side play a key role in the recovery of motor function. Immunofluorescence double labeling revealed that fiber scars are mainly distributed in the lesion core but glial scars are mostly localized in the rostral and caudal sites (the transition zone) (Fig.S2A). But, GV treatment increased the GFAP positive area largely in the centra of lesion and decreased fibrotic scar marked by laminin and fibronectin when comparing to SCC group at 4wpi (Fig.S2A-E, P<0.001 in Fig.S2C, P<0.01 in Fig.S2D, P<0.001 in Fig.S2E). Furthermore, quantities analyses showed that the relative translations of laminin and fibronectin were higher in the SCC group than that of GV-SCC and sham group (Fig.S2F-H). When the GV treatment was performed for 8 weeks and unapplied for 3 weeks, fibrotic scar tissue rich in fibronectin and laminin also formed at the lesion site after SCC. GV treatment group showed a significant reduction of fibronectin and of laminin-positive fibrotic scar tissue in the dorsal spinal cord compared to WT SCC group (Fig.3D-F, P<0.001 for both). Reduced level of laminin and fibronectin after GV revealed that the fiber scar decreased and an advantageous environment conducive to axonal regeneration or plasticity enhancement formed. While, in CPLX1^−/+^ SCC rats, fibronectin and of laminin-positive fibrotic scar tissue barely existed and their positive area were lower than GV-WT group (Fig.3D-F, P<0.001 for laminin (+) and P<0.01 for fibronectin(+)). These changes were also confirmed by WB results (Fig.3G-H).

### Moderate reduction of CPLX1 (CPLX1^−/+^) promotes axon regrowth through highly expressing the GAP43, and further increases synaptic plasticity and serotonergic innervation, which were all important for locomotion

Previously, increased serotonergic innervation strongly modulate the recovery of motor function after SCI (Mullner *et al.*, 2008). We then tested whether the scar-reducing effect also promotes axon regrowth of serotonergic spinal axons. Indeed, the CPLX1^−/+^ SCC rats showed increased density of 5-HT^+^ axons innervating the ventral horn caudal to the lesion compared to WT SCC animals (Fig.4A-B, P<0.05). And, GV administration also enhanced the 5-HT^+^ fibers compared with that of WT SCC models (Fig.4A-B, P<0.001). Same trends also happened in the longitudinal section of the WT, GV-WT and CPLX1^−/+^ SCC groups (left side of white dotted line showed the caudal side to the lesion core) (Fig.4A). Moreover, CPLX1^−/+^ promoted axon growth even when these neurons were in an environment that fully with the inhibitory molecules including Nogo-A, Semaphorin3A (Sema-3A) and CSPGs, abundant at the spinal cord lesion area (Fig.5A-B, P<0.001 for both of them). For the spinal cord neurons, moderate reduction of CPLX1 also played the same effect (Fig.S4). Which protein was attributed this effect? We detected that growth associated protein 43 (GAP43) was significantly up-regulated in CPLX1^−/+^ cortical neurons than in the WT group both at axon and neurosome (Fig.5,C-H, P<0.000 for axon analysis; P<0.05 for soma analysis).

**Figure 4.**
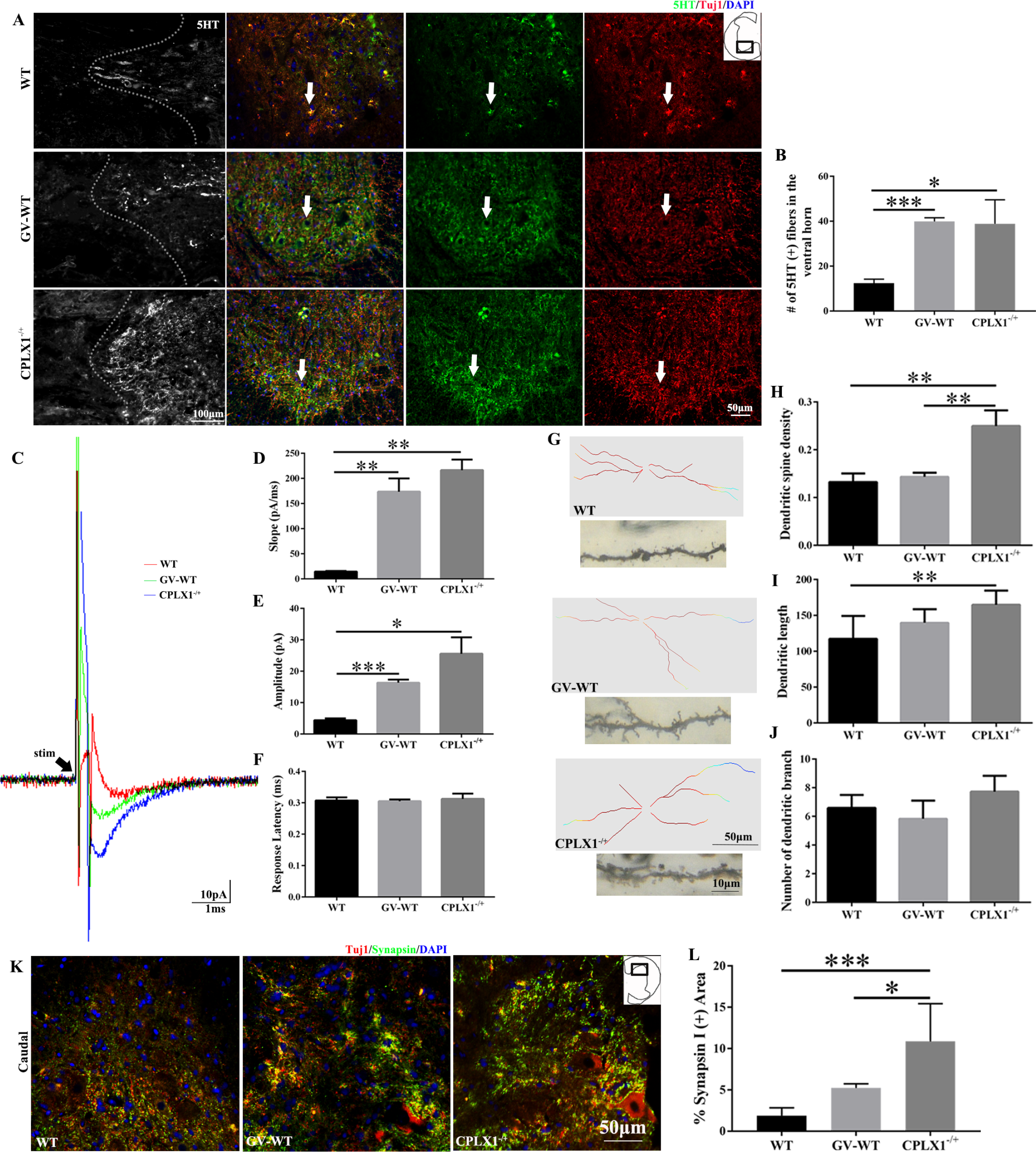
Moderate reduction of CPLX1 (CPLX1^−/+^) promotes regeneration of serotonergic spinal axons, enhances synaptic plasticity and increases the fEPSP caudal to the lesion even more than GV application. A Serotonin (5HT) immunolabeling (dashed line, lesion border) and longitudinal sections of the rat lumbar spinal cord after contusion injury. The second panel, double-staining of 5HT and Tuj1 in the transverse section of caudal spinal cord. The third and fourth panel, images of each marker visualizing serotonergic innervation of motor neurons (arrows). B The number of 5HT labeled (+) fibers caudal to a rat spinal cord, n = 5 animals per group. C Representative traces of averaged (30 trials) DC evoked fEPSCs from 4 weeks WT (red trace), GV-WT (green trace) and CPLX1^−/+^ (blue trace) SCC rat, n = 5 animals per group. D Quantitative data show the charge for evoked responses in three groups. E Group comparisons for dorsal column evoked peak amplitude. F Group comparisons for dorsal column evoked response latency. G Reconstructions of anterior horn motor neurons of caudal spinal cord in WT, GV-WT and CPLX1^−/+^ group show different dendritic patterns and dendritic spine density. H Quantitative data of dendritic spine density, n = 3 animals per group. I Quantitative data of dendritic length, n = 3 animals per group. J Quantitative data of dendritic branch, n = 3 animals per group. K Double immunolabeling of synapsin I (green) with Tuj1 (red). L Quantitative data of synapsin I positive area, n = 3 animals per group. Data information: Data were analyzed using one-way ANOVA. Values are plotted as means + SEM. *P < 0.05, **P < 0.01, ***P < 0.001.

**Figure 5.**
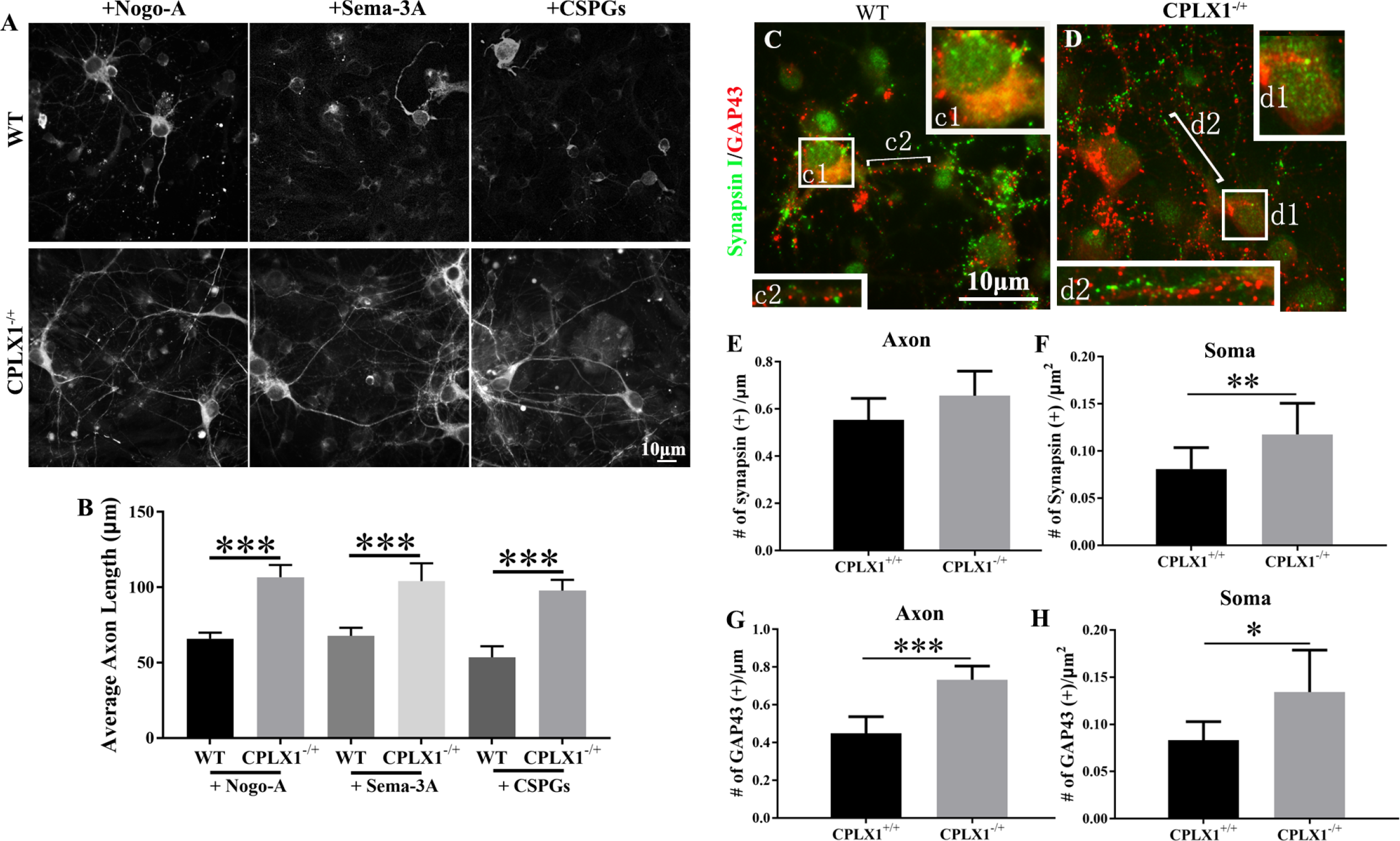
Moderate reduction of CPLX1 (CPLX1^−/+^) increases GAP43 expression and enhances axon elongation in cortical neurons. A Beta-3 tubulin (Tuj-1) immunolabeling of neurons growing on inhibitory substrates (Nogo-A; Sema-3A, Semaphorin-3A; CSPGs, chondroitin sulfate proteoglycans). B Neurite length of cortical neurons after 48 hours in WT and CPLX1^−/+^ genotype, n = 4 experiments. C, D Double immunolabeling of synapsin (green) with neuronal GAP43 (red). Boxed region in the low magnification image on the centra is shown at higher magnification immediately to the periphery. E, F Quantification of synapsin immunoreactivity at the axon and neuronal soma, n = 5 animals per group. G, H Quantification of GAP43 immunoreactivity at the axon and neuronal soma, n = 5 animals per group. Data information: Data were analyzed using Student’s t-test. Values are plotted as means + SEM. *P < 0.05, ***P < 0.001.

Further, we wondered that whether the scar-reducing effect also promotes spinal circuitry caudal to the lesion through increasing synaptic plasticity. To assess this, Golgi-stained motor neurons located throughout the caudal ventral horn was carried out for quantifying a fine details of neuron morphology. Results showed that the dendritic spine density of CPLX1^−/+^ SCC group was larger than GV and WT SCC groups (Fig.4G-H, P<0.01 for both comparisons). For the dendritic length, CPLX1^−/+^ SCC group was statistically longer than WT (Fig.4G, I, P<0.01). However, there was barely difference in the number of dendritic branch (Fig.4G, J). We next tested whether the increase in synaptic plasticity noted at the caudal to the injury epicenter was associated with synaptogenesis. Double stain of synapsin I and Tuj1 was used to quantify changes in presynaptic coverage on lumbar ventral horn motor neurons. There were significant main effects for both genotype (WT vs. CPLX1^−/+^, P<0.000) and different treatments (CPLX1^−/+^ vs. GV-WT, P<0.05) (Fig.4K-L; Fig.S5). Results of synapse in the lamellae 9 to 10 of ventral horns of spinal cords using transmission electron microscope also confirmed the synaptogenesis (Fig.S3). The synapse covered more in the groups of CPLX1^−/+^ SCC (P<0.0001) and GV-WT SCC (P<0.05) than the WT SCC group (Fig.S3). Besides, demyelination was more obvious in the SCC group but it was significantly improved when treating with GV or partial delection of CPLX1. To further study whether these changes could lead a local potential change might reflect the recovery of motor function of hindlimbs in the L2 spinal cord level, field excitatory postsynaptic potential (fEPSP) was measured. These data combined indicated that field EPSP were consistently evoked in all three groups. For the slope in the groups of GV-WT SCC and CPLX1^−/+^ SCC, they were higher than WT respectively (P<0.01), and in the amplitude of GV-WT (P<0.001) and CPLX1^−/+^ (P<0.05), they were higher than WT SCC group, while no obvious difference was observed in the response latency (Fig.4C-F).

### GV improves the neurological deficits via eIF5A1 regulating CPLX1

From the above, we proved that down-regulated CPLX1 in the effectiveness of GV treatment has a special contribution. We then assessed whether other molecule involved in this process. Protein sequence of CPLX1 is highly conserved in Rattus norvegicua, Mus musculus and Homo sapiens, and they all had the same PPG sequences which induce ribosomes become stalled resulting no full-length product produced (Fig.6A). However, previously reports demonstrated that eukaryotic eIF5A1 can rescue the stalled ribosomes (Doerfel *et al.*, 2013; Huter *et al.*, 2017). To verify whether eIF5A1 was the main reason for down-regulated CPLX1 in GV treatment, we firstly used the primary cultured cortical neurons to confirm the translated and functional regulation of eIF5A1 to CPLX1. Briefly, we cultured primary cortical neurons and transfected with HSVs carrying with eIF5A1-ORF or CPLX1-ORF, and siRNA targeting these two molecules. After overexpressing eIF5A1, the level of eIF5A1 increased, followed by the increasing of CPLX1 level, while eIF5A1 was down-regulated with the decreasing expression of CPLX1, and there was a reduction eIF5A1 when CPLX1 was overexpressed (Fig.6B-D). These observations suggested that eIF5A1 can positively regulate the gene translation of CPLX1. For the functional validation, FM1-43 dyes were used to label and then monitor synaptic vesicles, secretory granules and other endocytic structures in a variety of preparations. When stimulating with high potassium (K^+^), the intensity of FM1-43 fluorescence in CPLX1 siRNA and eIF5A1 siRNA group were higher than other groups (Fig.6E-F). However, the high signal phenomenon of eIF5A1 siRNA elicited by the low expression of CPLX1 was reversed after overexpressing CPLX1 (Fig. 6E-F). Afterwards, these results had described decreased exocytosis in neurons with lower-expressing of CPLX1. To further verify this, we examined potassium (K^+^) and calcium (Ca^2+^) fluxes in neurons using microelectrode ion flux estimation (MIFE). Before and after stimulating with high K^+^, K^+^ flux of CPLX1^−/+^ neurons was higher than that of WT neurons (P<0.05), while there was a significant reduction of the flux of CPLX1^−/+^-eIF5A1-ORF after overexpressing eIF5A1 (P<0.001) (Fig.6G-H). Accordingly, the flux of Ca^2+^ in the CPLX1^−/+^ group was much less than that of WT group (P<0.01), but after overexpressing eIF5A1, the flux of Ca^2+^ was increased obviously (P<0.05) (Fig.6I-J). Delection of CPLX1 (CPLX1^−/−^) also significantly decreased the efflux of Ca^2+^, which can be rescued by overexpressing CPLX1 but not eIF5A1 (Fig.6K). Therefore, rescue tests were proved that CPLX1 is functional regulated by eIF5A1 in vitro.

**Figure 6.**
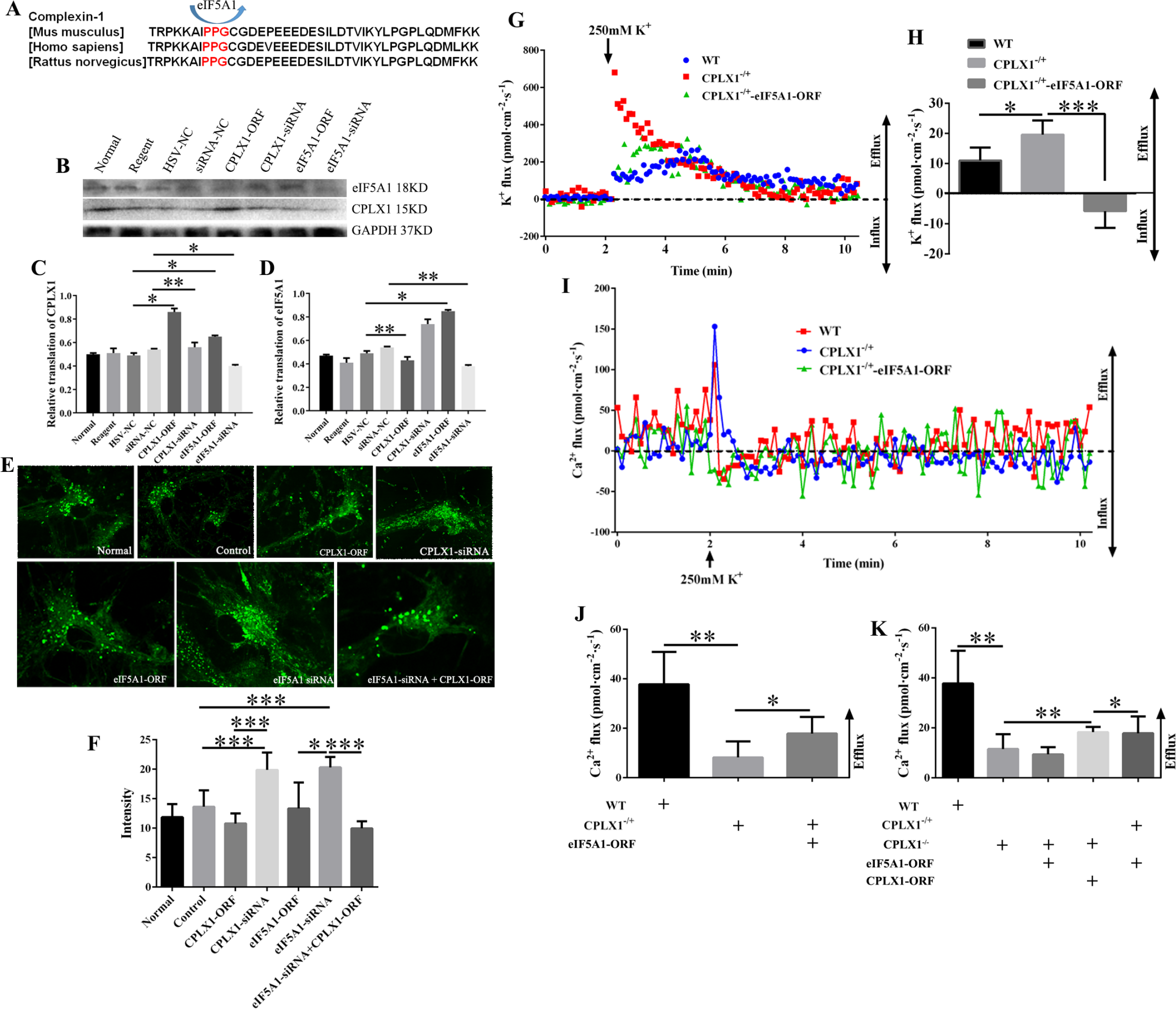
eIF5A1 promotes translation of CPLX1. A Amino acid sequence contained the PPG of CPLX1 in mouse, rat and human. B WB analysis of soluble proteins extracted from different treated spinal cord neurons. GAPDH served as a loading control. C Quantification of CPLX1 level among different groups. D Quantification of eIF5A1 level among different groups. E Exocytosis labeling by FM1-43 in cortical neurons processing differently. F Quantification of the fluorescence intensity among different groups. G K^+^ ion flux of primary cultured spinal cord neurons before and after high potassium treatment. H Baseline of K^+^ ion flux in WT or presence of either CPLX1^−/+^ or transfected with HSV-eIF5A1-ORF. I Ca^2+^ ion flux of primary cultured spinal cord neurons before and after high potassium treatment. J, K Baseline of Ca^2+^ ion flux among different groups. Data information: Data were analyzed using one-way ANOVA. Data are plotted as means ± SEM. *P < 0.05, **P < 0.05, ***P < 0.001.

To evaluate whether eIF5A1-dependent CPLX1 expression can affect the neurobehavioral recovery in SCC models, we used CPLX1^−/+^ rats to generate SCC model and HSV- eIF5A1-ORF was injected into the spinal cord lesion area of CPLX1^−/+^ rats. WB results confirmed that the content of CPLX1 was decreased as mentioned above in CPLX1^−/+^ SCC group compared to WT SCC group (P<0.001), but increased after over-expression of eIF5A1 in CPLX1^−/+^ SCC group (P<0.05) (Fig.7A). EMG recordings of these SCC models demonstrated that over-expression of eIF5A1 reverse the enhancement of amplitudes in CPLX1^−/+^ SCC group (P<0.05) (Fig.7B-C). Same trends also found in the MEP recordings in which the amplitude of CPLX1^−/+^ group was larger than WT group (P<0.01) and CPLX1^−/+^-eIF5A1-ORF group (P<0.05) (Fig.7D-E). And, the response latency in the CPLX1^−/+^ group was shorter when compared to the CPLX1^−/+^-eIF5A1-ORF group (P<0.01, Fig.7F). BBB scores analysis showed that CPLX1^−/+^-eIF5A1-ORF group had a lower BBB scores compared to CPLX1^−/+^ SCC models begin 4wpi, which revealing that over-expression of eIF5A1 in CPLX1^−/+^ SCC models reversed the improved neurobehavioral defects (Fig.7G). We finally investigated whether the motor function recovery and amelioration after GV treatment was duo to down-regulated CPLX1 through eIF5A1. HSV- eIF5A1-ORF was injected into the spinal cord lesion area of WT SCC models whose were given the GV treatment. Six weeks later, BBB scores of GV+ eIF5A1-ORF group was lower than that of GV-WT group (Fig.7H, P<0.05). The protein levels of eIF5A1 and CPLX1 were increased in the lesion area of GV+ eIF5A1-ORF group comparing with GV-SCC group (Fig.7I-K, P<0.01 for eIF5A1 detection and P<0.05 for CPLX1 detection).

**Figure 7.**
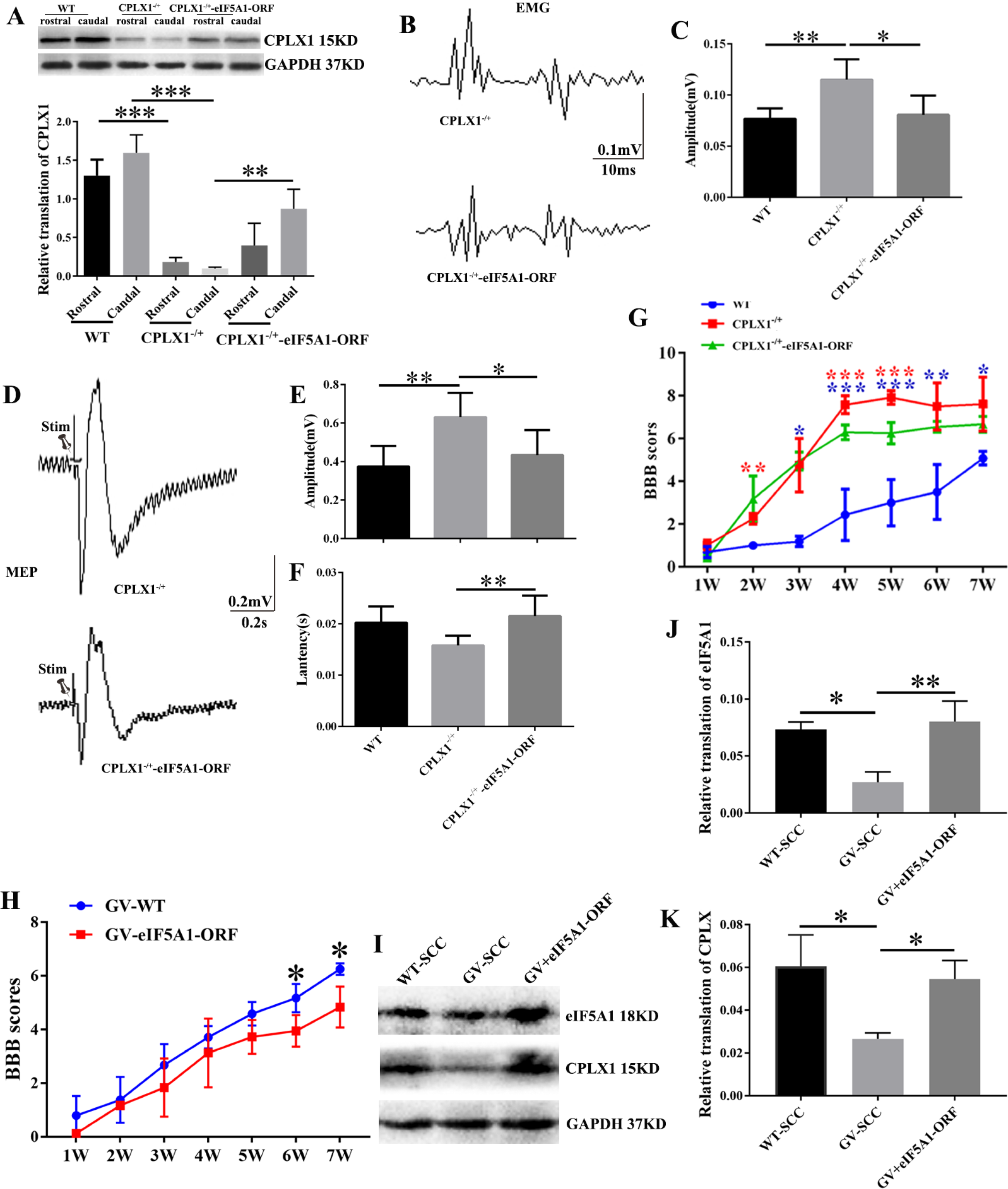
eIF5A1 stimulates the expression of CPLX1 in SCC models. A WB of CPLX1 in the caudal and rostral spinal cord extracts among different groups, n = 3 animals per group. B Representative EMG signals in different group, n=4 animals per group. C Quantification of amplitude among these indicated groups. D Representative MEP signals in different group, n=4 animals per group. E Quantification and statistical analyses of amplitude for the aforementioned experiments are shown. F Quantification of latency period for these groups. G, H BBB scores among these indicated group. I WB of CPLX1 and eIF5A1 in the injured spinal cord extracts of GV-WT and GV+eIF5A1-ORF groups, n = 3 animals per group. J, K Quantification of eIF5A1 and CPLX1 translation among different groups. Data information: Data were analyzed using one-way ANOVA. Data are plotted as means ± SEM. *P < 0.05, **P < 0.05, ***P < 0.001.

## Discussion

As well known, SCI induces widespread molecular and biochemical changes including altered mRNA and protein expression, axonal plasticity, and neuronal cell death, and there is no cure for drugs and methods. Nowadays, GV has been applied in treating SCI far and wide (Wong *et al.*, 2003; Mo *et al.*, 2016). However, the precise mechanisms of GV on neurogenesis are still largely unknown, which seriously limited the therapeutic effect. In this study, we were committed to find the underlying mechanism of GV treating SCI. Firstly, we verified the effect of GV by experimenting on the SCC rats. BBB scores showed that SCC rats get a remarkable motor recovery after GV. Besides, less formation of cavity, inflammatory cell infiltration, demyelination and increased synaptic plasticity were found after GV. Using protein chip, CPLX1was found to be the most possible target gene. It was observed that the level of CPLX1 was reduced after GV. In addition, we confirmed that lower CPLX1 expression promoted axon regeneration and synapse plasticity and motor function in vivo. Therefore, axon regeneration and synapse plasticity appear both contributing to the functional recovery, revealing a possible mechanism that GV promote function recovery in SCC models. Moreover, the observation of sequence PPG containing in CPLX1 requires eIF5A1 to rescue their translation providing further investigation. In this study, we attempted to find the underlying mechanism of GV via CPLX1 regulated by eIF5A1.

Previous studies have demonstrated that scarring represents a major barrier for axonal regrowth and that moderate inhibition of this process will enable axonal regrowth and improve functional recovery (Toy&Namgung, 2013). Here, fibrotic scar tissue rich in fibronectin and laminin forms at the lesion site after SCI, which forms a key impediment to regenerate axons, containing axon growth inhibitory factors, including chondroitin sulfate proteoglycans (CSPGs)(Ruschel *et al.*, 2013). Furthermore, reduced level of laminin and fibronectin after GV which assessed by immunofluorescence double labeling revealed that the fiber scar decreased and an advantageous environment conducive to axonal regeneration or plasticity enhancement formed. It was previously reported that secondary degeneration of neurons is reduced and axon regeneration is facilitated by GV (Wang *et al.*, 2016).

Besides, modulating the plastic changes at the spinal cord level play an important role in improving functional recovery (Asensio-Pinilla *et al.*, 2009). In this study, DTI was performed to track spinal cord neural regeneration progress. It was observed that the fiber signals filled the whole spinal cord structure in a continuously state after GV, which proved that GV could promote the axon regeneration (Feng&Zhang, 2014). In summary, we provide a strong rationale that GV could facilitate axon regeneration and improve recovery of motor function.

In order to investigate the underlying mechanism, results of protein chip show that the expression of CPLX1 decreased obviously after GV. CPLX1 is a highly charged protein that is essential for Ca^2+^-mediated neurotransmission that appears to act by interacting with and regulating the SNAREs (Chen *et al.*, 2007). To date, CPLX1 levels are differentially expressed in many psychiatric and neurodegenerative disorders (Zhou *et al.*, 2017). Studies have shown that CPLX1 is specifically existed in the nervous system and pancreatic B cells, and expressed in neurons, microglia, and astrocytes, especially mainly expressed in the synaptic structure-rich region (McMahon *et al.*, 1995). In recent years, it has been found that CPLX1 is a molecule with two-way regulation function, which exerts both positive and negative effects on vesicle exocytosis, facilitating synchronous neurotransmission while inhibiting spontaneous fusion events (Wragg *et al.*, 2013; Zdanowicz *et al.*, 2017). Moreover, more studies have reported that CPLX1 are crucially involved in neurological development and neurotransmitter release and the availability and importance of CPLX1 in nerve repair have been expounded in many studies (Ugurel *et al.*, 2016). Given the important role of CPLX1 in nervous system, we want to ascertain whether GV effectively improves motor function of SCC rats through down-regulation of CPLX1. So, CPXL1^−/+^ rats with lower expression of CPLX1 were selected in subsequent experiments to investigate the role of CPXL1 in SCC damage. Here, CPLX1^−/+^ rats had a better motor recovery as revealed by higher BBB scores and increased EMG and MEP signals. The objectivity of SEP and MEP recording ensures that lower expression of CPLX1 enhances the electrophysiological conduction through the injured spinal cord and the excitability of the sciatic nerve, which further confirm the BBB scores obtained (Iyer *et al.*, 2010).

Up to now, one of the important strategies for treating SCC is to promote axon regeneration in the epicenter (Vigneswara *et al.*, 2012). We then investigated whether GV treatment promotes regrowth of descending axons, which mediate voluntary motor movement, important for locomotion (Dias *et al.*, 2018). Increased serotonergic (5-HT) axons play a key role in activating and modulating lumbar motor circuitry after GV treatment (Freria et al., 2017). Previously studies report that 5-HT axons have an intrinsic ability to sprout and grow after SCI, bypassing the epicenter and repopulating gray matter caudal to the injury site, which has been implicated in spontaneous recovery of motor function (Leech et al., 2014; Freria et al., 2017). Here, the density 5-HT^+^ axons caudal to the epicenter enhanced in SCC rats either deficient in CPLX1 (CPLX1^−/+^) or performed the GV treatment. Moreover, the regrowth of 5-HT^+^ axons repopulate lumber spinal cord gray matter showing a topographically appropriate pattern around the anterior motor neurons.

GV also promotes synaptic plasticity via regulating CPLX1. Numerous evidences show that the dendritic spines can change shape, size, and number following various injuries (Tan&Waxman, 2012; Deng *et al.*, 2016). After SCI, there is an acute reduction in dendritic number in these survived neurons with rapid dendritic atrophy. Meanwhile, spontaneous dendritic plasticity could reflect a compensatory response of the spinal cord to the functional deficits caused by the injury (Horch *et al.*, 2011; Deng *et al.*, 2016). Increased dendritic spine density observed in neurons have entered into a more plastic state, which indicate availability to form new synapses (Alvarez&Sabatini, 2007). Indeed, data in the current report indicate that low expression of CPLX1 can increase the number of neurons, dendritic branch and dendritic length in the caudal site to the epicenter. Other reports demonstrate that dendritic plasticity is formed through actin cytoskeleton reorganization which responds to glutamate release regulate downstream signals such as PKA, PKC, MAPK and Rac1 through activating AMPA and NMDA receptors (Tan&Waxman, 2012). As CPLX1 plays an important role in the process of neurotransmitters release (Kummel *et al.*, 2011), thus enhanced the synaptic plasticity of neurons may result from the low expression of CPLX1 clamping chaotic release of neurotransmitters caused by SCI. Field potential demonstrated that the regenerated descending axons across the lesion and activated the intraspinal circuits below the lesion contributing to the majority of functional recovery (Deng *et al.*, 2013). Here, myelination of survived axons increased in the group of low expression of CPXL1 and GV treatment, suggesting that these axons could conduct action potentials. These evidences in our models demonstrated that regenerated axons and enhanced synaptic plasticity caused an increased fEPSP, promoting the recovery of motor function in rats with SCC, suggesting that is beneficial for the recovery of spinal cord injury, which may be related to the involvement of CPLX1 in synaptic plasticity. Taken together, it can be drawn the conclusion that CPXL1 affects the neuronal synaptic plasticity participating in GV treatment of spinal cord injury by regulating neurotransmitter release.

In particular, CPLX1 is a highly conserved protein with the sequences PPG, which can stall the translation. Moreover, many researches had confirmed that eIF5A1 can rescue the translation of sequences PPG (Gutierrez *et al.*, 2013; Nguyen *et al.*, 2015), so we made an assumption that eIF5A1 can regulate CPLX1 in the response to the neurobehavioral deficits in SCC. In this study, we proved that eIF5A1 can positively regulate the expression of CPLX1 protein. The regulatory verification of SCC function recovery *in vitro* and release of neuronal vesicle neurotransmitters *in vivo* was carried out, indicating that overexpression of CPLX1 can reverse neuronal vesicle release dysfunction caused by eIF5A1 interference. On the other hand, overexpression of eIF5A1 can reverse the recovery of motor function and better fiber remodeling caused by low expression of CPLX1. Meanwhile, overexpression of eIF5A1 increased the metabolic activity of the upper and lesion tissues in the WT rats with SCC. It was found that overexpression of eIF5A1 increased the expression of CPLX1 in the rostral and caudal to lesion sites in the CPLX1^−/+^ SCC model.

## Materials and Methods

### Animals and materials

Adult female Sprague-Dawley rats (200-250g) were obtained from animal center of Sichuan University. All procedures were followed by international, national and institutional guidelines and were approved by local authorities (Sichuan Medical Experimental Animal Care Commission #2016098A). Animals were housed in a comfortable and clean condition under a 12/12-h dark-light cycle following SCC experiment, and food was available ad libitum. Moreover, their bladders were manually massaged three times daily. Complexin1 knock-out rats were provided by Institute of laboratory animal sciences, CAMS&PUMC. HSV-vector was obtained from Brain VTA.

### Rat model of Spinal Cord Contusion (SCC)

After deeply anesthetized with a ketamine (80 mg/kg, i.p.)/xylazine (10 mg/kg, i.p.) mixture, rats were fixed in a prone position. A midline skin incision was made between the area of T10 and L2, and then paravertebral muscles were separated. After exposing the spinal cord, the vertebrae were stabilized with clamps and the rats were suffered a moderate (75 kdyn) mid-thoracic (T10) contusion SCI (PCI3000 Precision Cortical Impactor, Hatteras Instruments, Inc). While in the sham group, the rats only exposed the spinal cord without contusion after anesthetized. After impaction, the surgery incision was sutured and rats were hydrated with 2 ml of saline (intraperitoneal injection) and were given prophylactic antibiotic (0.1ml cefotaxime sodium, i.p.) for 3 days. Manual bladder expression was performed 3 times a day until recovery of micturition reflex.

### Basso, Beattie, and Bresnahan (BBB) Score

The recovery of motor function in hind limbs was evaluated using a Basso, Beattie, and Bresnahan open field locomotion rating scoring system (BBB score). BBB score was performed on weekly after injury. In brief, this scale used for assessing hindlimb function includes evaluation standards ranging from a score of 0, indicating no spontaneous movement, to a maximum score of 21, with an increasing score indicating the use of individual joints, coordinated joint movement, coordinated limb movement, weight-bearing, and other functions. Three researchers “blinded” to rat treatment status performed 5min tests on all animals.

### MRI and DTI

Each rat was anesthetized with 3% isflurone before MRI scan, and anesthesia was maintained during the scan by continuous administration of mixture gas (2% isflurone −O_2_/N_2_ (30:70)). During MRI, the temperature, heart rate and breathe of rats was monitored periodically. The images were captured with T2WI sequence (TR/TE = 2000/15ms, NAverages=4, FOV=50mm×50mm, RareFactor=8), and each scanning period lasted 4 min 16 s. DTI was performed after MRI. Echo planar imaging sequence was applied with 15 gradient directions (b=1000 s/mm2, echo time=32.2ms, Segments=8, RePET/CTition time=2000ms, FOV=35mm × 35 mm, matrix =128 × 128), and scanning is lasted for 42 min 40 s. After obtaining the image, each data was corrected using DTI studio (Jiang *et al.*, 2006). Then the Diffusion Toolkit software was used to perform white matter fiber tracking on the corrected DTI data and display the image using Trackvis (http://www.trackvis.org).

### DTI data processing

DTI scans were processed and analyzed by means of dedicated MRICRON (https://www.softpedia.com/get/Science-CAD/MRIcron.shtml), DTI Studio (http://cmrm.med.jhmi.edu/), Trackvis. Firstly, we used MRICRON to transpose Dicym to Nifti. The Automatic Image Registeration was conducted for all DW scans within DTI Studio. The deformation field is calculated in the same way for each of all DW scans and applied accordingly. Afterwards, Deuffusion Toolkit was performed to track matter fiber. After processing, three vertical eigenvalues are derived from each pixel to calculate b0 and FA. RY image was captured and FA as well parameters of fiber was got using Trackvis. The direction of the eigenvector associated with the largest eigenvalue is set to the main direction of the local neural fiber. The background removal threshold of 0.10 was set to exclude non-normal voxels and any significant noise; the smoothing of the interpolating fiber was set to 20% and the minimum fiber length was set to 1 cm for continuous fiber reconstruction. The FA values in the ROI at the rostral, lesion and caudal sites were extracted and used for statistical comparison.

### MEP

After being anaesthetized by 3% isoflurane, the limbs of rats were abducted and fixed on a board by cloth bands. Successive stimulation was given at musculi hippicus, and the recorded contraction of the target muscle was taken at the stimulating intensity between 3.5 and 12 mA. A pulse train of 5 pulses was used, with pulse width 100 s, intra-pulse period of 50ms, and inter-train frequency of 0.5 Hz. The MEP signal was recorded for 500ms after the initiation of each pulse-train, but only the portion of the MEP signal located within 50ms of the final stimulus pulse was analyzed. These stimulation frequencies were chosen through reference to papers by (Schlag *et al.*, 2001). MEP was recorded from each of the limbs using sub-dermal needle electrode pairs. The grand average of all time-locked sweeps was taken and then was used for all further analysis.

### EMG

After anesthetized by 3% isflurone, the upper back of each rat was shaved and cleaned and a small skin incision was made in the motor area of cerebral cortex to allow placement of the transmitter. The wire electrodes of the transmitter were tunneled subcutaneously to the right hindlimb by separating the skin from the muscle layer using blunt dissection. EMG recordings were made from rats to track the changes that occurred in reflexes over time.

### Field potential

Horizontal spinal cord slices were prepared according to the previous reports (Rank *et al.*, 2015). Briefly, Horizontal slices (300 μm thick) were cut using a vibrating microtome (VT1200 S, Leica Biosystems, Germany) submerged in oxygenated, ice-cold, sucrose-substituted artificial cerebrospinal fluid (sACSF) (in mM): 250 sucrose, 2.5 KCl, 1 CaCl_2_, 6 MgCl_2_, 25 NaHCO_3_, 11 glucose and 1 NaH_2_PO_4_ (PH 7.4). Then the slice was immediately transferred to a recovery bath and perfused with oxygenated containing ACSF (in mM): 118 NaCl, 2.5 KCl, 2.5 CaCl_2_, 1 MgCl_2_, 25 NaHCO_3_, 1 NaH_2_PO_4_, 11 glucose and pH 7.4 with NaOH. The slice was positioned in the bath, and secured under a custom-made net. The sections were held around 270 mV by injecting a hyperpolarisation current, and the spike was elicited by a 500 pA depolarising ramp current for 100ms. Recordings were made at 37°C using an Axopatch 200B amplifier (Molecular Devices, Sunnyvale, CA, USA). Data were collected (sampled at 50 kHz, filtered between 2 and 10 kHz) and analysed on a computer using pCLAMP 10 software (Molecular Devices Corp., Sunnyvale, CA 94089 USA).

### Tissue Harvest

Animals performed for WB were deeply anesthetized. After rats were decapitated, the spinal cord tissues rostral and caudal to the lesion site were fetched quickly and were placed into phosphate buffered saline (PBS) at 4°C for following experiments. For immunofluorescent staining, the samples were collected after intracardiac perfusion with 50 ml physiological saline followed with 4% paraformaldehyde. Additionally, the tissue samples for Golgi staining should be collected instantly after anaesthetizing rats. After removal, the surface blood was quickly washed away with H_2_O_2_, and the samples were soaked in the immersion liquid.

### HE Staining

Firstly, frozen sections were washed by distilled water for 2 min, and then were stained using hematoxylin and eosin. After rinsing in tap water for 10 min to wash off any excess staining solution followed by distilled water once more and 0.95% ethanol for 5 seconds, sections were counterstained with Eosin, dehydrated, cleared with xylene and mounted with neutral gum. The stained sections were observed using OlyVIA slice scanner to detect morphologic changes.

### Immunocytochemistry

After animals were perfused with 4% paraformaldehyde solution, spinal cord segments were harvested and dehydrated by 30 % sucrose overnight. For in vivo staining, sagittal 1cm sections using freezing microtome (Leica CM1900, Germany) encompassing regions both rostral and caudal to the lesion. Mounted sections were washed three times with PBS followed by blocking in 5% normal goat serum(NGS) and 0.1% Triton X-100 in PBS. After blocking, sections were incubated in primary antibody diluted in 2% NGS overnight at 4 °C. Primary antibodies used were mouse anti-GFAP (1:200, #MAB 3402, Chemicon), mouse anti-Tuj1 (1:200, #MAB1195, R&D systems), rabbit anti-laminin (1:100, #ab11575, Abcam), rabbit anti-fibronectin (1:200, #ab2413, Abcam) and rabbit anti-5HT (1:50, #ab10385, abcam), and rabbit anti-synapsin I (1:500, #SAB4502904, Sigma-Aldrich). The next day, the sections were washed extensively with PBS and incubated in the appropriate secondary antibody conjugated to AlexaFluor 488 and 594 (1:400, Molecular Probes) overnight. After extensive washing, the sections were stained with DAPI, coverslipped and viewed with a confocal microscope (Zeiss, Germany). Pixel intensity was measured on images taken on a standard fluorescent microscope (Leica) with a uniform exposure setting and analysed using ImageJ.

### Western Blotting

Protein was extracted from frozen spinal cord tissue samples (200 mg), then lysed and homogenized in RIPA lysis buffer containing 2 % of cocktail pill (Roche). All samples were centrifuged at 12000 ×g for 10 min at 4°C. Then the total supernatant protein was collected and its concentration was determined by BCA protein assay (Thermo ScientificTM, #23225). After separating samples containing 80 μg of protein on 10% sodium dodecyl sulfate polyacrylamide gel electrophoresis, the separated proteins were transferred onto PVDF membranes, and the membranes were blocked for 1 h with TBST buffer containing 5% skim milk. Membranes were incubated overnight at 4 °C with the following primary antibodies, rabbit anti-CPLX1 (1:1000, #17700, SCT); rabbit anti-Laminin (1:1000, #ab11575, Abcam); rabbit anti-Fibronectin (1:5000, #ab2413, Abcam), then were incubated for 2 h at room temperature with Horseradish peroxidase-coupled secondary antibodies (1:5000, Abcam). GAPDH was used as a loading control (1:50000, #AC033, Abclonal). All samples were visualized using ECL detection reagents (Beyotime, China). Quantitative densitometric analysis was processed using Image J software.

### Golgi Staining

Golgi Staining was performed using the FD Fast Golgi Staining Kit (#PK401, FD NeuroTechnologies, Inc.) according to the protocol. The spinal cord tissues were soaked in the mixture of solution A and B for 3weeks, and then transfer into solution C for 72 h in dark place. Then the fixed tissues were sectioned at a thickness of 100um at −22°C using a thermostated microtome (CM1860, Leica). Tissues were stained followed by rinsing with double distilled water for 4 mins, three times. Later, sections were placed in the mixture (1x D, 1x E and 2x distilled water) for 10 min. After repeating washing with distilled water twice, sections were dehydrated in 50%, 75%, and 95% and absolute ethanol for 4 minutes, soaked in xylene and mounted with neutral gum. The image was taken using an OlyVIA 2.9 slice scanner (OLYMPUS).

### Primary culture of spinal cord neurons

Neonatal rats (less than 24 hours) were used for primary cell culture. Animals were decapitated at the base of the foramen magnum after sterilization, vertebral columns were separated along the sagittal suture. Then, using the microscissors to open the spinal canal, the spinal cord was harvested, minced, and isolated by 0.25% trypsinase for 10min at 37°C following by complete medium (DMEM (high glucose) with 10% BSA) to neutralize the effect of trypsinase. The cells were collected by centrifugation at 1,000 rpm for 10 min, resuspended by complete medium, and plated in 6-well plates (Corning, USA) at the density of 10^5^ cells/ml. After incubation at 37 °C with 5 % CO2 for 4 h, the culture medium previously used was replaced by Neurobasal added with B27.

### Lenti-virus Transection in spinal cord neurons

6 days later, cells were transfected by HSV virus containing different open reading frame (ORF) with MOI=1. Then, images were acquired using Leica AF6000 at 3 days after transfection.. Five fields were used for measuring soma size and number as well as neurite length of cortical neurons by using Leica DMI6000B (LAS AF system) and mean value was calculated. Then, the size, number, and neurite length were measured, respectively.

### Synaptic bouton activity (FM1-43)

After clearing the supernatant in the cultured primary neuron, primary neuron was washed twice with PBS. Cells were incubated for 10 minutes in a 2 ml low-K^+^ buffer, then were incubated for 5 minutes in high-K^+^ buffer containing 10 mM FM1-43 dye, followed by washing with a low-K^+^ buffer to remove the surface-bound dye. Later, cells were stimulated for 5 minutes in a high-K^+^ medium at 37°C. The extra liquid was removed and formaldehyde was used to fix. Images were captured using a Leica TCS SP5 microscope equipped with a HCX PL APO 63X 1.4 numerical aperture oil immersion objective.

### Electron microscopic analysis

Sections were cut from the tissues of spinal cord using an ultramicrotome (Leica EM FC7), and then placed on formva-coated grids. Ultrathin sections were examined in a FEI Tecnai G2 F20 transmission electron microscope at 80 kV. Images were acquired by camera (EagleTM4K CCD).

### Ion-selective flux measurements

Complete experimental procedure of microelectrode ion flux estimation (MIFF) has been processed by a modified procedures according to the previously reports (Shabala *et al.*, 2010). Cortical neurons for the MIFE detections were cultured for 6 days at 5× 10^5^ cells/well on coverslips coated with poly-L-lysine. Cells were washed in ASCF (0.5 mM CaCl_2_, 5 mM KCl, 52 mM NaCl, 26.2 mM NaHCO_3_, 0.9 mM NaH_2_PO_4_, 0.5 mM MgCl2, 5 mM D-Glucose, 2 mM HEPES, PH 7.4) and then placed into a measuring chamber immerged into 2ml ASCF. Measurements were performed after three days post transfected with different HSV virus. The K^+^ and Ca2+ fluxes were recorded for 5 min prior to the addition of a high-K^+^ solution (250 mM KCl, 0.2 mM CaCl_2_, 52 mM NaCl, 26.2 mM NaHCO_3_, 0.9 mM NaH_2_PO_4_, 0.5 mM MgCl2, 5 mM D-Glucose, 2 mM HEPES, PH 7.4), and recordings continued for 15 min once the testing liquid (0.2 mM CaCl_2_, 0.2 mM KCl, 52 mM NaCl, 26.2 mM NaHCO_3_, 0.9 mM NaH_2_PO_4_, 0.5 mM MgCl2, 5 mM D-Glucose, 2 mM HEPES, PH 7.4) was immediately added to exchange the high-K^+^ solution. For recording the K^+^ and Ca^2+^ fluxes, microelectrodes were filled with 100 mM KCl and 100 mM CaCl_2_, respectively. Data was collected at a rate of 10 samples/sec and averaged over 6 second intervals. Net ion fluxes (nmolm^−2^s^−1^) were analysed and calculated using MIFE software.

### Statistical Analysis

All data in the experiment are presented as mean ± standard deviation. Measurement data were statistically analyzed utilizing SPSS 21.0 software (SPSS, Inc., Chicago, USA). Differences between groups were compared by two sample t-tests. P<0.05 was regarded as statistically significant.

## Acknowledgements

This work was supported by grant of National Natural Science Foundation of China (NSFC) (NO.81471268 and NO.81271358).

## Author contributions

YX and THW conceived and supervised the project. JL, XMZ and LZ designed and performed the experiments together with YJ, CYL, YYW and LLX analyzed data. YX wrote the manuscript with contributions and approval of all the other authors.

## Conflict of interest

The authors declare that they have no conflict of interest.

## The paper explained

### Problem

GV treatment can effectively improve the motor function of SCC models through reducing scar and promoting axon regeneration, which is driven by an intense signals. But key molecules governing this process have not been identified so far.

### Results

Using a CRISPR/Cas9 knockout approach, CPLX1^−/+^ rats were used to assess the role of CPLX1 in GV treatment. Additionally, eIF5A1 stimulate the translation of CPLX1 with PPG sequence, we attempt to uncover whether eIF5A1 play a role in the GV treatment. In fact, GV can reduce scar and promote axon regeneration after SCC. Decreased CPLX1 improved the microenvironment of injured area via reducing the components of fibrotic scar and further enhanced the synaptic plasticity, which benefit the regeneration of axons. And eIF5A1 could regulate the expression of CPLX1 in the process of GV treatment.

### Impact

Our experimental data have demonstrated that CPLX1 functions as a neurogenesis gene in promoting synapse plasticity and axon regeneration. These findings evaluated a detailed mechanism of GV for SCC treatment, providing a new insight that GV exerts its role via eIF5A1 regulating CPLX1.

---

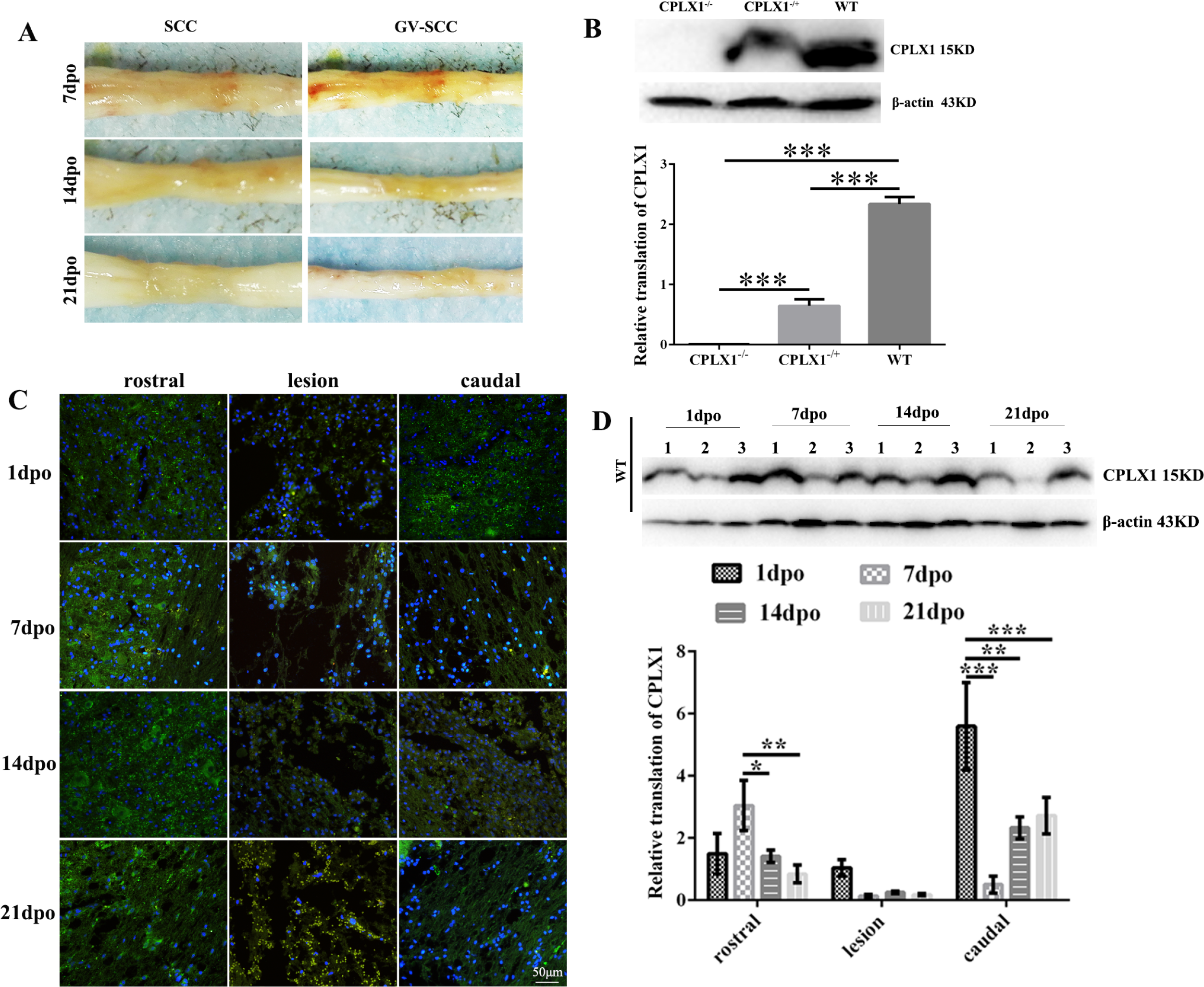

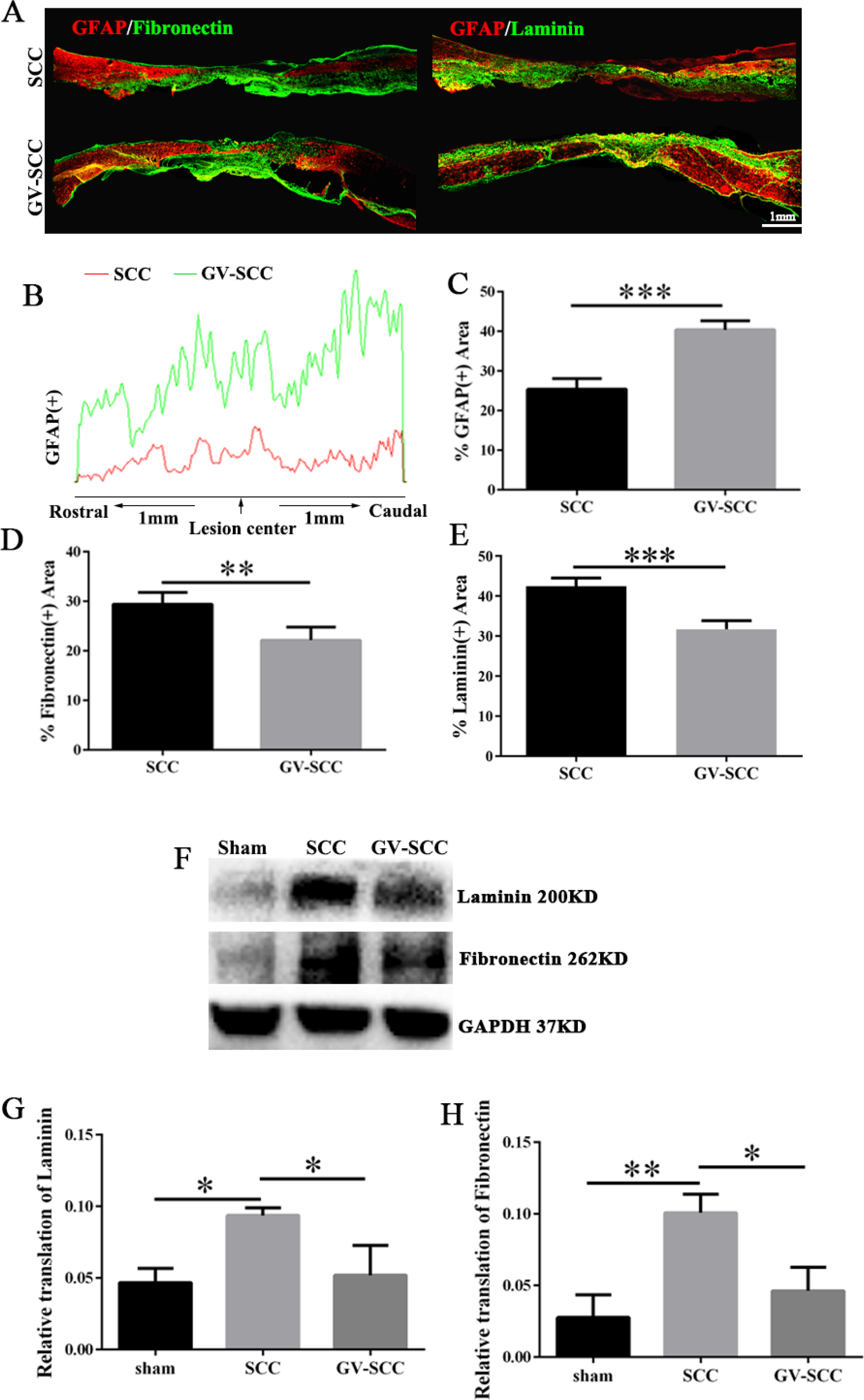

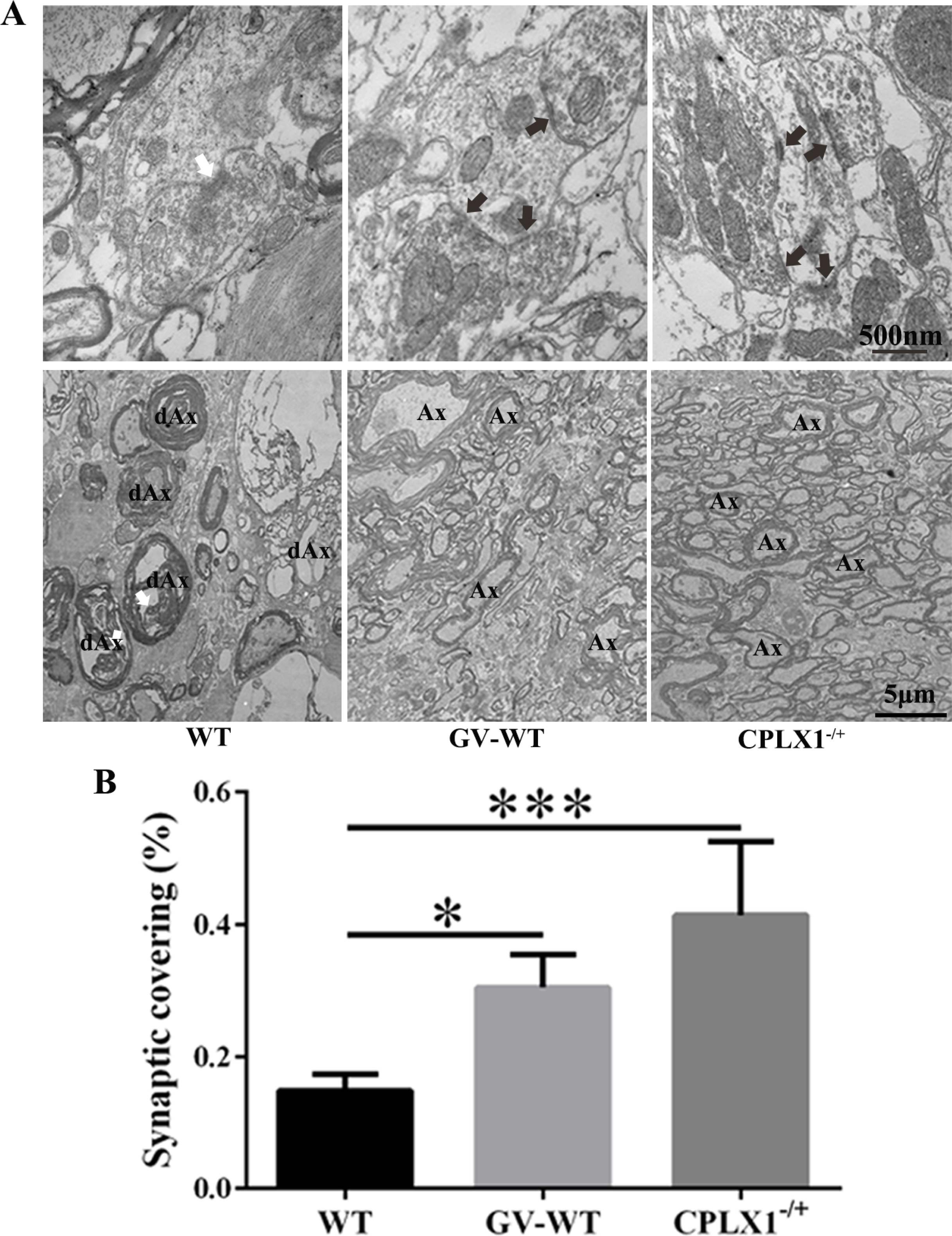

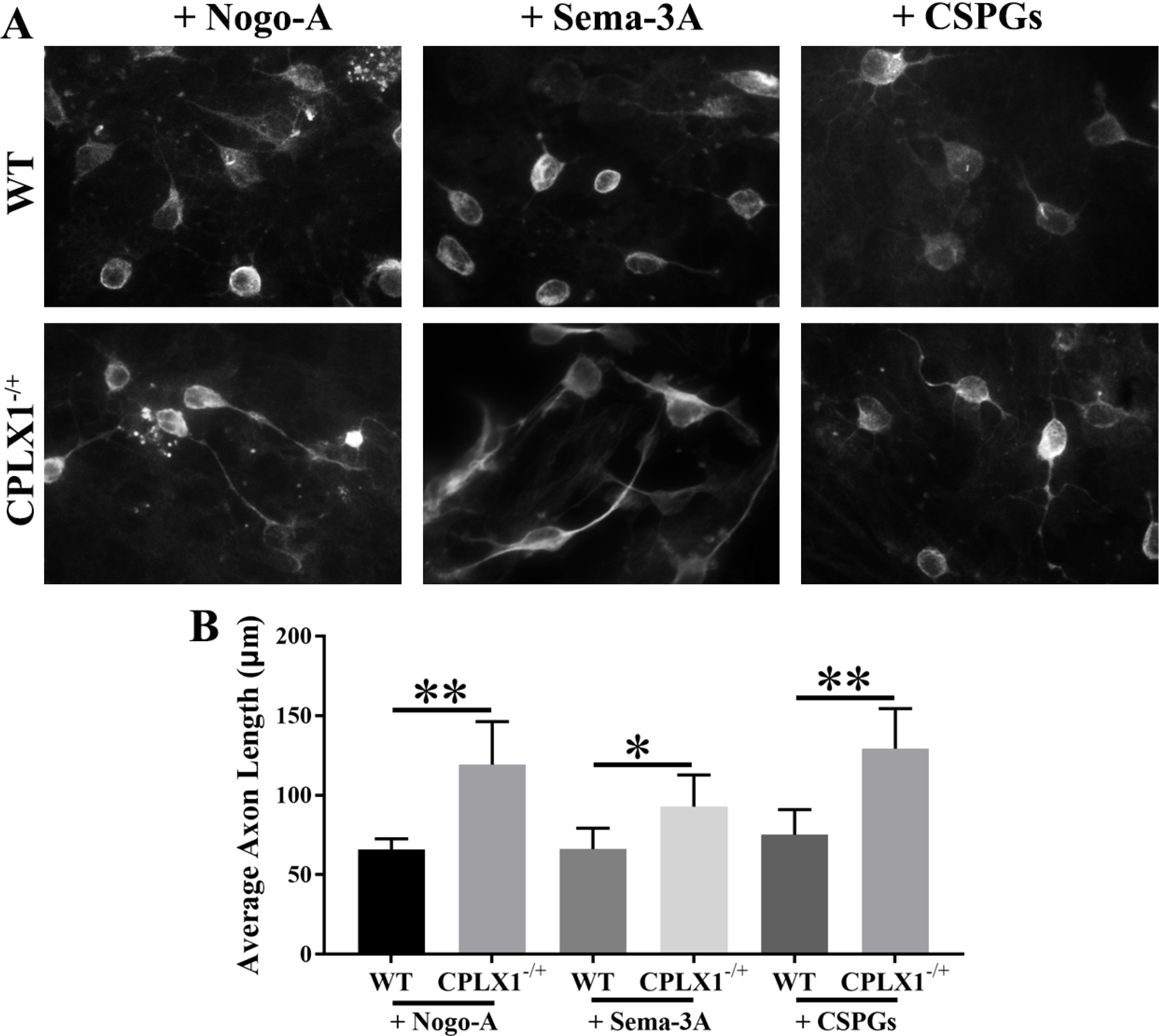

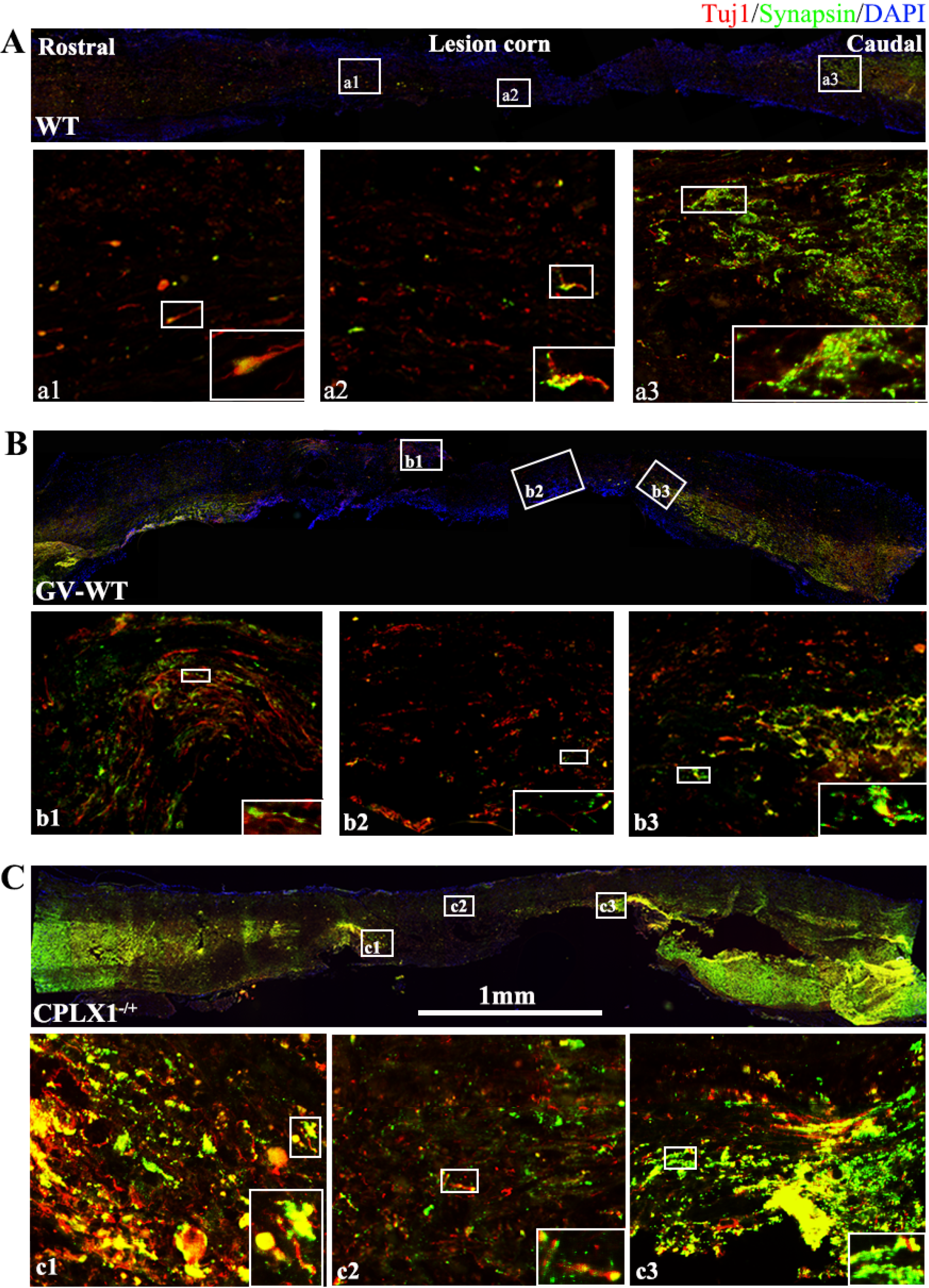

## References

Alvarez VA, Sabatini BL (2007) Anatomical and physiological plasticity of dendritic spines. Annual Review of Neuroscience 30: 79–97

Asboth L, Friedli L, Beauparlant J, Martinez-Gonzalez C, Anil S, Rey E, Baud L, Pidpruzhnykova G, Anderson MA, Shkorbatova P, Batti L, Pages S, Kreider J, Schneider BL, Barraud Q, Courtine G (2018) Cortico-reticulo-spinal circuit reorganization enables functional recovery after severe spinal cord contusion. Nat Neurosci 21: 576–588

Asensio-Pinilla E, Udina E, Jaramillo J, Navarro X (2009) Electrical stimulation combined with exercise increase axonal regeneration after peripheral nerve injury. Exp Neurol 219: 258–265

Chen B, Li Y, Yu B, Zhang Z, Brommer B, Williams PR, Liu Y, Hegarty SV, Zhou S, Zhu J, Guo H, Lu Y, Zhang Y, Gu X, He Z (2018) Reactivation of Dormant Relay Pathways in Injured Spinal Cord by KCC2 Manipulations. Cell 174: 521–535.e513.

Chen J, Qi JG, Zhang W, Zhou X, Meng QS, Zhang WM, Wang XY, Wang TH (2007) Electro-acupuncture induced NGF, BDNF and NT-3 expression in spared L6 dorsal root ganglion in cats subjected to removal of adjacent ganglia. Neurosci Res 59: 399–405

Deng LX, Deng P, Ruan YW, Xu ZC, Liu NK, Wen XJ, Smith GM, Xu XM (2013) A Novel Growth-Promoting Pathway Formed by GDNF-Overexpressing Schwann Cells Promotes Propriospinal Axonal Regeneration, Synapse Formation, and Partial Recovery of Function after Spinal Cord Injury. Journal of Neuroscience 33: 5655–5667

Deng LX, Ruan YW, Chen C, Frye CC, Xiong WH, Jin XM, Jones K, Sengelaub D, Xu XM (2016) Characterization of dendritic morphology and neurotransmitter phenotype of thoracic descending propriospinal neurons after complete spinal cord transection and GDNF treatment. Exp Neurol 277: 103–114

Dias DO, Kim H, Holl D, Werne Solnestam B, Lundeberg J, Carlen M, Goritz C, Frisen J (2018) Reducing Pericyte-Derived Scarring Promotes Recovery after Spinal Cord Injury. Cell 173: 153–165 e122

Ding Y, Yan Q, Ruan JW, Zhang YQ, Li WJ, Zhang YJ, Li Y, Dong H, Zeng YS (2009) Electro-acupuncture promotes survival, differentiation of the bone marrow mesenchymal stem cells as well as functional recovery in the spinal cord-transected rats. BMC Neurosci 10: 35

Doerfel LK, Wohlgemuth I, Kothe C, Peske F, Urlaub H, Rodnina MV (2013) EF-P Is Essential for Rapid Synthesis of Proteins Containing Consecutive Proline Residues. Science 339: 85–88

Feng R, Zhang F (2014) The neuroprotective effect of electro-acupuncture against ischemic stroke in animal model: a review. Afr J Tradit Complement Altern Med 11: 25–29

Freeman W, Morton AJ (2004) Regional and progressive changes in brain expression of complexin II in a mouse transgenic for the Huntington’s disease mutation. Brain Res Bull 63: 45–55

Freria CM, Hall JCE, Wei P, Guan Z, McTigue DM, Popovich PG (2017) Deletion of the Fractalkine Receptor, CX3CR1, Improves Endogenous Repair, Axon Sprouting, and Synaptogenesis after Spinal Cord Injury in Mice. Journal of Neuroscience 37: 3568–3587

Glynn D, Reim K, Brose N, Morton AJ (2007) Depletion of Complexin II does not affect disease progression in a mouse model of Huntington’s disease (HD); support for role for complexin II in behavioural pathology in a mouse model of HD. Brain Res Bull 72: 108–120

Gutierrez E, Shin BS, Woolstenhulme CJ, Kim JR, Saini P, Buskirk AR, Dever TE (2013) eIF5A promotes translation of polyproline motifs. Mol Cell 51: 35–45

Hazell AS, Wang C (2005) Downregulation of complexin I and complexin II in the medial thalamus is blocked by N-acetylcysteine in experimental Wernicke’s encephalopathy. J Neurosci Res 79: 200–207

Horch HW, Sheldon E, Cutting CC, Williams CR, Riker DM, Peckler HR, Sangal RB (2011) Bilateral Consequences of Chronic Unilateral Deafferentation in the Auditory System of the Cricket Gryllus bimaculatus. Developmental Neuroscience 33: 21–37

Huter P, Arenz S, Bock LV, Graf M, Frister JO, Heuer A, Peil L, Starosta AL, Wohlgemuth I, Peske F, Novacek J, Berninghausen O, Grubmuller H, Tenson T, Beckmann R, Rodnina MV, Vaiana AC, Wilson DN (2017) Structural Basis for Polyproline-Mediated Ribosome Stalling and Rescue by the Translation Elongation Factor EF-P. Mol Cell 68: 515–+

Iyer S, Maybhate A, Presacco A, All AH (2010) Multi-limb acquisition of motor evoked potentials and its application in spinal cord injury. J Neurosci Methods 193: 210–216

Jiang HY, van Zijl PCM, Kim J, Pearlson GD, Mori S (2006) DtiStudio: Resource program for diffusion tensor computation and fiber bundle tracking. Computer Methods and Programs in Biomedicine 81: 106–116

Kummel D, Krishnakumar SS, Radoff DT, Li F, Giraudo CG, Pincet F, Rothman JE, Reinisch KM (2011) Complexin cross-links prefusion SNAREs into a zigzag array. Nature Structural & Molecular Biology 18: 927–U1603

Leech KA, Kinnaird CR, Hornby TG (2014) Effects of Serotonergic Medications on Locomotor Performance in Humans with Incomplete Spinal Cord Injury. Journal of Neurotrauma 31: 1334–1342

Li M, Peng J, Song Y, Liang H, Mei Y, Fang Y (2012) Electro-acupuncture combined with transcranial magnetic stimulation improves learning and memory function of rats with cerebral infarction by inhibiting neuron cell apoptosis. J Huazhong Univ Sci Technolog Med Sci 32: 746–749

Li WJ, Li SM, Ding Y, He B, Keegan J, Dong H, Ruan JW, Zeng YS (2012) Electro-acupuncture upregulates CGRP expression after rat spinal cord transection. Neurochem Int 61: 1397–1403

Liu J, Wu Y (2017) Electro-acupuncture-modulated miR-214 prevents neuronal apoptosis by targeting Bax and inhibits sodium channel Nav1.3 expression in rats after spinal cord injury. Biomed Pharmacother 89: 1125–1135

Liu Z, Ding Y, Zeng YS (2011) A new combined therapeutic strategy of governor vessel electro-acupuncture and adult stem cell transplantation promotes the recovery of injured spinal cord. Curr Med Chem 18: 5165–5171

Maeda Y, Kim H, Kettner N, Kim J, Cina S, Malatesta C, Gerber J, McManus C, Ong-Sutherland R, Mezzacappa P, Libby A, Mawla I, Morse LR, Kaptchuk TJ, Audette J, Napadow V (2017) Rewiring the primary somatosensory cortex in carpal tunnel syndrome with acupuncture. Brain 140: 914–927

McMahon HT, Missler M, Li C, Sudhof TC (1995) Complexins: cytosolic proteins that regulate SNAP receptor function. Cell 83: 111–119

Mo YP, Yao HJ, Lv W, Song LY, Song HT, Yuan XC, Mao YQ, Jing QK, Shi SH, Li ZG (2016) Effects of Electroacupuncture at Governor Vessel Acupoints on Neurotrophin-3 in Rats with Experimental Spinal Cord Injury. Neural Plasticity

Moraud EM, Capogrosso M, Formento E, Wenger N, DiGiovanna J, Courtine G, Micera S (2016) Mechanisms Underlying the Neuromodulation of Spinal Circuits for Correcting Gait and Balance Deficits after Spinal Cord Injury. Neuron 89: 814–828

Mullner A, Gonzenbach RR, Weinmann O, Schnell L, Liebscher T, Schwab ME (2008) Lamina-specific restoration of serotonergic projections after Nogo-A antibody treatment of spinal cord injury in rats. Eur J Neurosci 27: 326–333

Nguyen S, Leija C, Kinch L, Regmi S, Li Q, Grishin NV, Phillips MA (2015) Deoxyhypusine Modification of Eukaryotic Translation Initiation Factor 5A (eIF5A) Is Essential for Trypanosoma brucei Growth and for Expression of Polyprolyl-containing Proteins. J Biol Chem 290: 19987–19998

Nie L, Xia J, Li H, Zhang Z, Yang Y, Huang X, He Z, Liu J, Yang X (2017) Ginsenoside Rg1 Ameliorates Behavioral Abnormalities and Modulates the Hippocampal Proteomic Change in Triple Transgenic Mice of Alzheimer’s Disease. Oxid Med Cell Longev 2017: 6473506

Rank MM, Flynn JR, Battistuzzo CR, Galea MP, Callister R, Callister RJ (2015) Functional changes in deep dorsal horn interneurons following spinal cord injury are enhanced with different durations of exercise training. Journal of Physiology-London 593: 331–345

Ruschel J, Bradke F (2018) Systemic administration of epothilone D improves functional recovery of walking after rat spinal cord contusion injury. Exp Neurol 306: 243–249

Ruschel J, Hellal F, Flynn C, Dupraz S, Blesch A, Weidner N, Bunge MB, Bixby JL, Bradke F (2013) Systemic Administration of Epothilone B Promotes Axon Regeneration and Functional Recovery after Spinal Cord Injury. Molecular Biology of the Cell 24

Schlag MG, Hopf R, Redl H (2001) Serial recording of sensory, corticomotor, and brainstem-derived motor evoked potentials in the rat. Somatosensory and Motor Research 18: 106–116

Shabala L, Howells C, West AK, Chung RS (2010) Prolonged Ab treatment leads to impairment in the ability of primary cortical neurons to maintain K+ and Ca2+ homeostasis. Molecular Neurodegeneration 5

Tan AM, Waxman SG (2012) Spinal cord injury, dendritic spine remodeling, and spinal memory mechanisms. Exp Neurol 235: 142–151

Toy D, Namgung U (2013) Role of glial cells in axonal regeneration. Exp Neurobiol 22: 68–76

Ugurel E, Sehitoglu E, Tuzun E, Kurtuncu M, Coban A, Vural B (2016) increased complexin-1 and decreased miR-185 expression levels in Behcet’s disease with and without neurological involvement. Neurol Sci 37: 411–416

Vigneswara V, Kundi S, Ahmed Z (2012) Receptor tyrosine kinases: molecular switches regulating CNS axon regeneration. J Signal Transduct 2012: 361721

Wang X, Duffy P, McGee AW, Hasan O, Gould G, Tu N, Harel NY, Huang Y, Carson RE, Weinzimmer D, Ropchan J, Benowitz LI, Cafferty WB, Strittmatter SM (2011) Recovery from chronic spinal cord contusion after Nogo receptor intervention. Ann Neurol 70: 805–821

Wang Y, Tang Q, Zhu L, Huang R, Huang L, Koleini M, Zou D (2016) Effects of Treatment of Treadmill Combined with Electro-Acupuncture on Tibia Bone Mass and Substance PExpression of Rabbits with Sciatic Nerve Injury. PLoS One 11: e0164652

Wong AMK, Leong CP, Su TY, Yu SW, Tsai WC, Chen CPC (2003) Clinical trial of acupuncture for patients with spinal cord injuries. American Journal of Physical Medicine & Rehabilitation 82: 21–27

Wragg RT, Snead D, Dong Y, Ramlall TF, Menon I, Bai J, Eliezer D, Dittman JS (2013) Synaptic vesicles position complexin to block spontaneous fusion. Neuron 77: 323–334

Zdanowicz R, Kreutzberger A, Liang B, Kiessling V, Tamm LK, Cafiso DS (2017) Complexin Binding to Membranes and Acceptor t-SNAREs Explains Its Clamping Effect on Fusion. Biophys J 113: 1235–1250

Zhang K, Liu Z, Li G, Lai BQ, Qin LN, Ding Y, Ruan JW, Zhang SX, Zeng YS (2014) Electro-acupuncture promotes the survival and differentiation of transplanted bone marrow mesenchymal stem cells pre-induced with neurotrophin-3 and retinoic acid in gelatin sponge scaffold after rat spinal cord transection. Stem Cell Rev 10: 612–625

Zhou Q, Zhou P, Wang AL, Wu D, Zhao M, Sudhof TC, Brunger AT (2017) The primed SNARE-complexin-synaptotagmin complex for neuronal exocytosis. Nature 548: 420–425

